# scRL: Utilizing Reinforcement Learning to Evaluate Fate Decisions in Single-Cell Data

**DOI:** 10.1101/2024.07.04.602019

**Authors:** Zeyu Fu, Chunlin Chen, Song Wang, Junping Wang, Shilei Chen

**Affiliations:** State Key Laboratory of Trauma and Chemical Poisoning, Institute of Combined Injury, Chongqing Engineering Research Center for Nanomedicine, College of Preventive Medicine, Army Medical University, 400038, Chongqing, China; Department of Rehabilitation Medicine, The First Affiliated Hospital, Sun Yat-sen University, Guangzhou 510080, China

**Keywords:** Reinforcement learning, Fate decisions, Single cell

## Abstract

The rapid development of single-cell sequencing offers an unparalleled opportunity to delineate the heterogeneity of individual cells. However, current methods struggle to pinpoint the states of cell fate decisions. In this study, we introduce a novel approach called Single-cell Reinforcement Learning (scRL), which integrates reinforcement learning into single-cell data analysis using an actor-critic architecture. Among existing dimensionality reduction methods, we identified one with the best interpretability. Based on this latent space and combined with our scRL algorithm, we assessed the intensity of fate decisions at the single-cell level. Extensive evaluations demonstrate that scRL outperforms existing techniques, as well as their variants and alternative approaches, in assessing cell fate decisions. Moreover, scRL offers an alternative method for evaluating the intensity of cell lineage differentiation which shows competitive interpretability as well. The superiority of scRL in assessing fate decisions is confirmed across several types of single-cell datasets.

## Introduction

Single-cell sequencing has emerged as a transformative technology for the exploration of cellular heterogeneity at an unprecedented resolution of individual cells^1–3^. A key advancement related to single-cell sequencing is pseudotime analysis, as it can unravel the sequential order of cellular events and the transition states in a virtual timeline. Cell state transitions manifest as branching differentiation trajectories in the two-dimensional reduced space of the data^4,5^. Embedding techniques such as Principal Component Analysis (PCA), t-distributed Stochastic Neighbor Embedding (t-SNE), and Uniform Manifold Approximation and Projection (UMAP) have proven exceptionally useful for single-cell data analysis ^6–8^. Particularly, UMAP can adeptly preserve both local and global structures, so that advanced applications like PAGA, Monocle3, Slingshot, and CAPITAL leverage UMAP to enhance representations of the differentiation manifold for trajectory inference^9–12^.

Despite being widely served for revealing the differentiation trajectories and temporal ordering, the current methodologies fall short in addressing where and when cell fate decisions occur during differentiation, as well as in providing a framework to evaluate these decisions. In fact, cellular differentiation is not merely a linear pathway but a complex landscape where progenitors become lineage-restricted at various points and times influenced by a myriad of factors^13^. This insight challenges the traditional hierarchical model of differentiation and highlights the importance of identifying specific decision points where cells commit to a particular lineage or fate. Given that single-cell analysis has highlighted the dynamic nature of the differentiation process, it should consider both spatial and temporal dimensions when studying cell fate decisions.

Reinforcement Learning (RL) is a branch of machine learning with distinct characteristics that an employed agent learns to make decisions by interacting with the environment. Normally, the agent seeks to achieve a goal by performing actions and receiving feedback in the form of rewards or penalties. In recent years, RL has excelled in addressing sequential decision-making challenges and been applied in multiple areas such as robot control, gaming intelligence, and navigation^14–16^. The power of reinforcement learning mainly lies in its adaptability and its ability to iteratively improve its understanding of the process of a certain event. Given that the process of cellular differentiation within the single-cell landscape is inherently sequential and decision-centric, it is achievable to utilize RL for single-cell sequencing analysis of cellular differentiation. By leveraging its ability to explore new paths and exploit known pathways efficiently, RL is expected to pinpoint the key differentiation nodes and pathways, to facilitate a deep decoding of cell fate decisions. In this study, we introduce an approach termed scRL by integrating reinforcement learning into single-cell data analysis to resolve the challenges in decoding lineage and cell fate decisions. Briefly, scRL utilizes a grid world created from a UMAP two-dimensional embedding of high-dimensional data, followed by an actor-critic architecture to optimize differentiation strategies and assess fate decision strengths. The effectiveness of scRL is demonstrated through its ability to closely align pseudotime with distance trends in the two-dimensional manifold and to correlate lineage potential with pseudotime trends. Being applied to the single-cell datasets of acute myeloid leukemia (AML), human hematopoiesis and mouse endocrinogenesis, scRL is shown to have prominent capability to accurately pinpoint fate decision states and pre-expression states in complex multilineage systems. The practicality and superiority of scRL are further confirmed with forecasting and dynamic gene perturbation of lineage and cell fate decisions during hematopoietic stem cells (HSCs) differentiation under physio pathological conditions.

## Results

### The architecture of scRL

As shown in Figure 1, the process of scRL begins by constructing a grid embedding with a two-dimensional reduction space of the cell data managed by t-SNE, PCA, or UMAP, to put up a structured environment for navigating cell differentiation. In the implementation, edge detection algorithms identifying boundary points on this grid, which help in defining a fully connected graph structure, to ensure connectivity across the grid from the most primitive to the most mature cell states. For the reinforcement learning setup, scRL utilizes an actor-critic architecture^17^ working in the grid environment filled with predefined reward signals. These rewards are configured to either decay or increase exponentially with calculated pseudotime, reflecting the maturity of cell states. The value network evaluates the potential returns from each state, while the policy network decides on the next action, guiding the agent through the grid to explore different cell fates.

**Figure 1:**
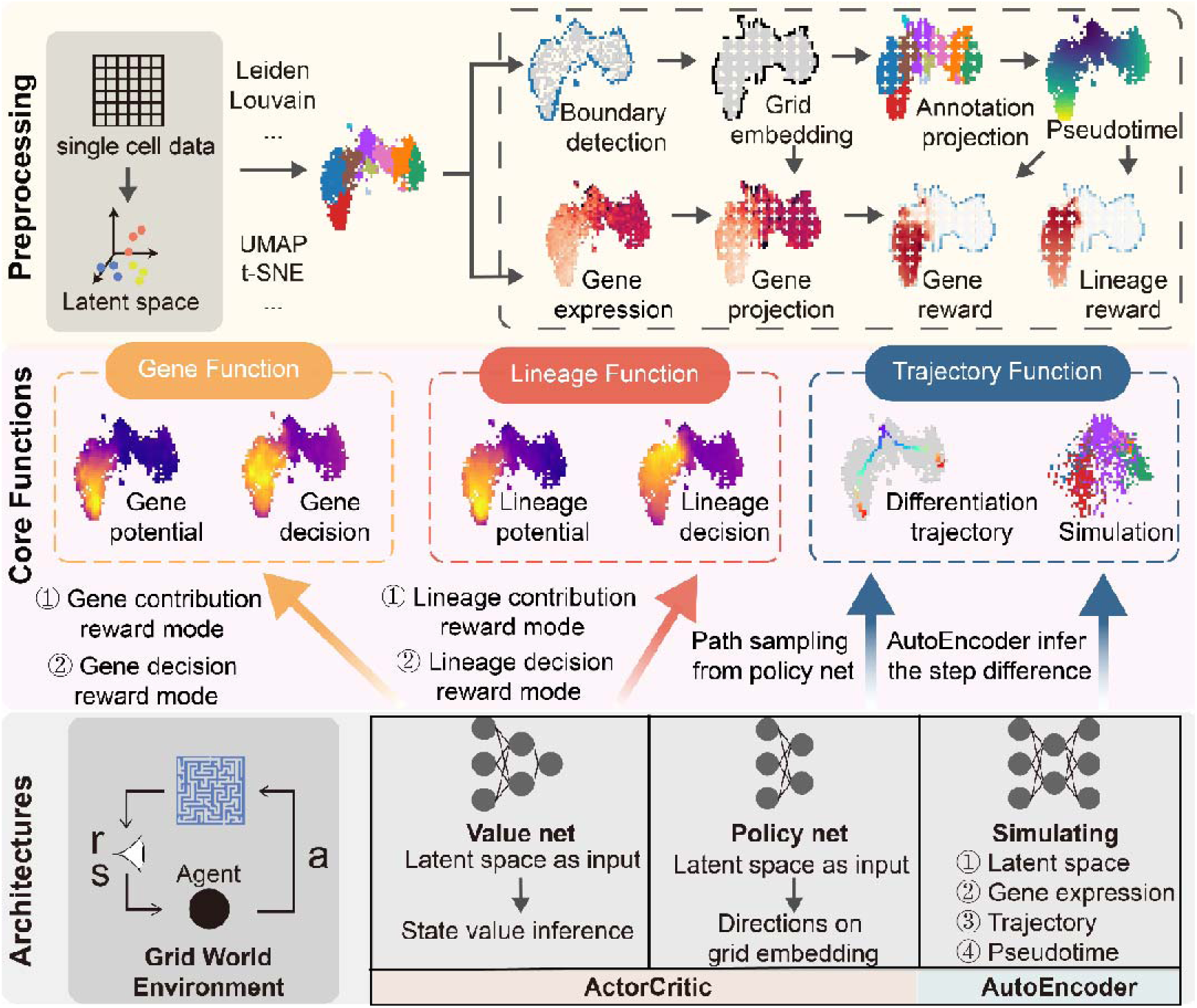
The architecture of scRL. The top panel of the figure outlines the preprocessing steps. It commences with Principal Component Analysis, progressing to dimensionality reduction and clustering techniques rooted in the principal component space. Through an edge detection algorithm, a grid embedding representing the original space emerges within the derived two-dimensional framework. Next, subpopulation information is mapped onto this grid embedding, and a pseudotime is computed from a designated starting subpopulation. Parallel to this, projecting gene expression data from the initial space onto the grid embedding gives rise to a gene reward environment. These dual reward environments, governed by their dynamic relationship with pseudotime shifts, yield a decision mode that diminishes along the pseudotime and a contribution mode that escalates in tandem. The central portion of the figure showcases scRL’s core functional modules, primarily segmented into three primary categories: gene, lineage, and trajectory modules. The gene module incorporates gene potential, derived from the contribution reward mode, alongside the gene decision value, emanating from the decision reward mode. The lineage module encompasses lineage potential from the contribution reward mode and lineage decision value from its decision counterpart. Lastly, the trajectory module encompasses both the differentiation trajectory and its simulation. The figure’s lower panel reveals the underlying structure of scRL. It commences with a pattern of the interplay between a reinforcement learning agent and its surroundings. This prelude segues into the Actor-Critic algorithm architecture. Here, the value net processes the principal component space, outputting the state value for each node. The policy net, operating on the same input space, selects differentiation strategies. An AutoEncoder is then utilized to simulate differentiation, processing the latent space from the preceding step, gene expression data, differentiation pathway, and pseudotime, subsequently generating predictions for the ensuing transition.

The core functionalities of scRL are divided into three modules: gene, lineage, and trajectory. The gene module uses gene expression data to train networks that predict gene decision strength and potential. The lineage module focuses on subgroup classification to determine fate decision strength and contribution strength of cell differentiation. The trajectory module samples and analyzes paths through the grid, representing differentiation trajectories.

Finally, an autoencoder^18^ is trained to predict changes in cell features like gene expression and pseudo-time, based on the differentiation trajectories identified by the policy network. This setup allows scRL to effectively map and predict cell differentiation processes, providing insights into the dynamics of cell fate decisions without the need to delve deeply into the technical specifics of the method.

### Exploring a Highly Interpretable Cellular Dimensionality Reduction Space

To investigate the mechanisms underlying cell fate decision, it is crucial to obtain a dimensionality reduction space that effectively captures cell differentiation trajectories. Therefore, we sought a single-cell dimensionality reduction method with excellent interpretability. In recent years, machine learning and deep learning techniques have been widely applied to the dimensionality reduction of single-cell data. In this study, we compared deep learning-based methods such as single-cell Variational Inference (scVI), Linear scVI, and Latent Dirichlet Allocation (LDA) with machine learning-based methods including PCA, Independent Component Analysis (ICA), Factor Analysis (FA), Non-negative Matrix Factorization (NMF), and Diffusion Map.

Using single-cell data from hematopoietic and pancreatic tissues, we applied techniques under varying numbers of highly variable genes and latent space dimensions to perform dimensionality reduction. We computed the maximal value labels of the latent components obtained from each method and evaluated them using discriminative metrics such as Adjusted Rand Index (ARI), Normalized Mutual Information (NMI), Average Silhouette Width (ASW), Calinski-Harabasz index, and Davies-Bouldin index. The LDA technique demonstrated significant superiority, achieving the highest overall score (Fig. 2A-B). Further analysis of the Silhouette scores corresponding to each latent component of each technique showed that LDA consistently had stable positive scores (Fig. 2C), indicating that the latent components obtained by LDA have the best interpretability among all the techniques evaluated. Additionally, we observed the distribution of LDA’s latent component intensities within their original subgroup labels, which exhibited strong subgroup specificity (Fig. 2D).

**Figure 2:**
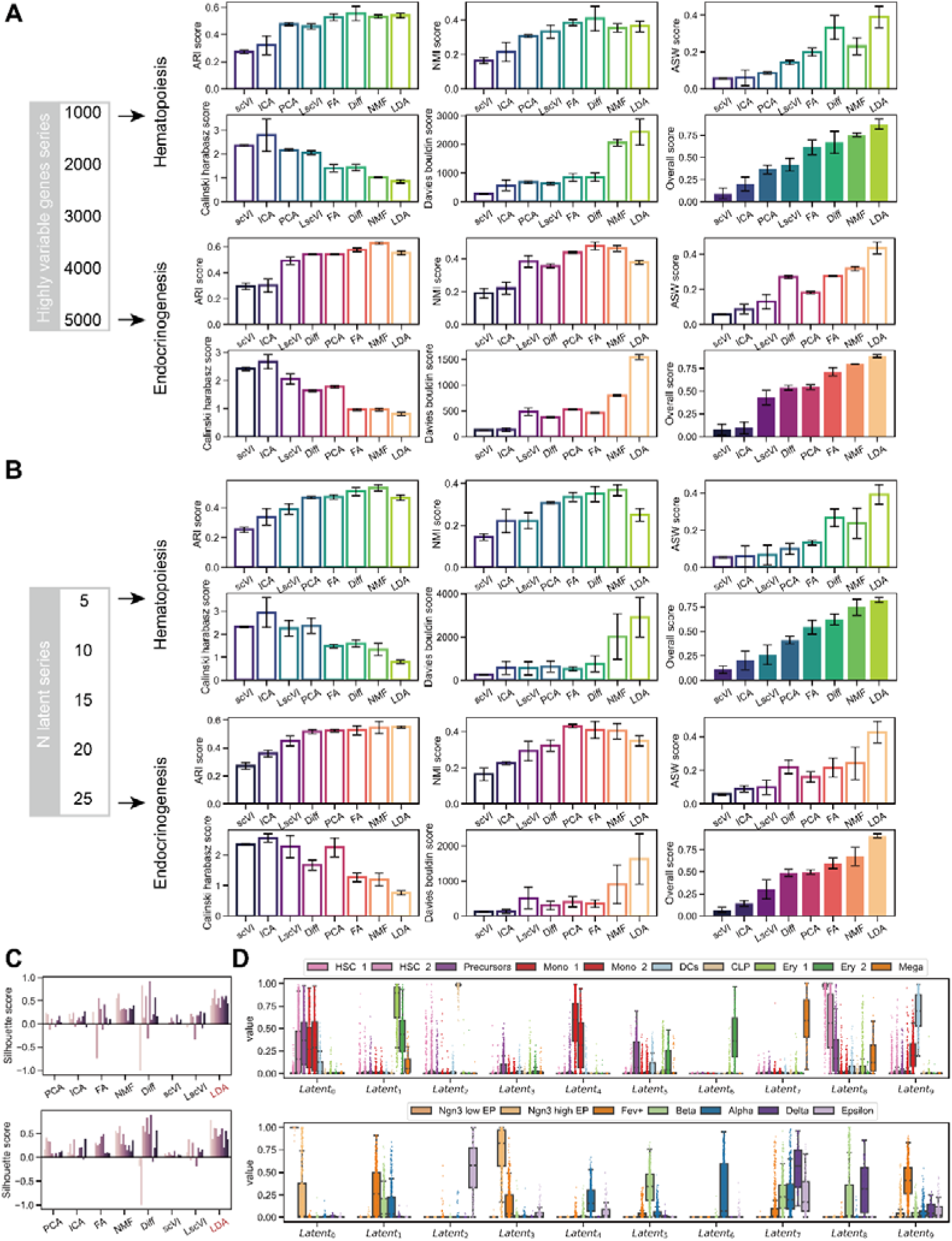
LDA Exhibits Superior Interpretability in Comparative Analysis of Various Single-Cell Dimensionality Reduction Methods. (A-B) Comparison of interpretability metrics such as ARI, NMI, ASW, Calinski, and Davies-Bouldin for dimensionality reduction methods including scVI, ICA, PCA, LscVI, FA, Diffusion Map, NMF, and LDA. The comparison is conducted under sequences of highly variable genes set at 1000, 2000, 3000, 4000, 5000, and latent space components set at 5, 10, 15, 20, 25. (C) Comparison of the average silhouette score for each subgroup obtained by various methods. (D) Intensity of latent components obtained by LDA across different subgroups.

### scRL Achieves Highly Interpretable Fate Decision Intensity

The intensity of lineage differentiation can be represented by lineage intensity, which is key to analyzing fate decision intensity. We attempted to use the intensities of each component in the interpretable LDA latent space as lineage intensities and projected them onto the corresponding two-dimensional UMAP embeddings. In the hematopoietic and pancreatic datasets, these lineage intensities corresponded to differentiation branches on the two-dimensional differentiation trajectories (Fig. 3A-B). To analyze fate decision intensity, scRL utilizes these trajectory branches, detects their edges, and constructs a grid embedding. Within this grid embedding, we can employ reinforcement learning to address the sequential decision-making problem of cell differentiation. By inferring the value of decisions, we obtain the corresponding intensity with which a cell chooses a certain fate at a given point, referred to as fate decision intensity (Fig. 3C).

**Figure 3:**
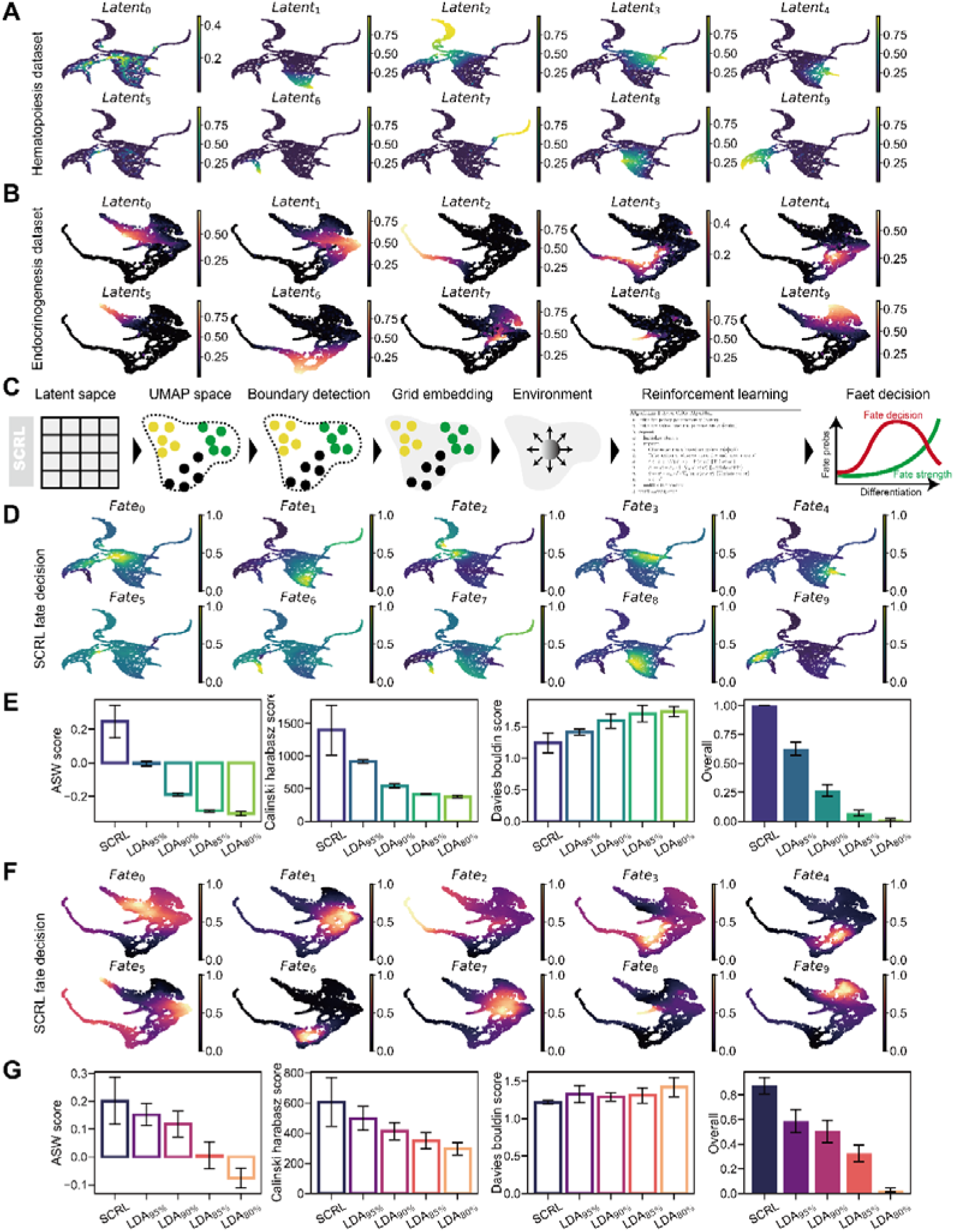
scRL Demonstrates Superior Interpretability in Fate Decision Intensity Analysis Compared to Using LDA Alone. (A-B) Projection of the intensity of latent space components obtained by LDA onto a 2D UMAP embedding in hematopoietic and endocrinogenesis datasets. (C) Workflow of scRL. (D, F) Projection of fate decision intensity obtained by scRL onto UMAP in hematopoietic and endocrinogenesis datasets. (E, G) Comparison of scRL and LDA at 95%, 90%, 85%, and 80% percentile intensities in hematopoietic and endocrinogenesis datasets using ASW, Calinski-Harabasz, and Davies-Bouldin metrics.

Projecting the fate decision intensity obtained by scRL onto the original two-dimensional differentiation trajectories, we found that the peak regions of fate decision intensity preceded those of lineage intensity along the trajectories. Therefore, to easily obtain the fate decision intensity of a particular lineage, an intuitive alternative is to advance the peak of the lineage intensity. We then compared the interpretability of the 95th, 90th, 85th, and 80th percentile intensities of the lineage intensities obtained by LDA, serving as alternative measures for fate decision intensity, with the fate decision intensity derived from scRL. Under the ASW, Calinski-Harabasz, and Davies-Bouldin metrics, scRL demonstrated superior interpretability compared to the alternative provided by the LDA technique in both test datasets (Fig. 3D-E).

### scRL’s Fate Decision Intensity Outperforms Existing Methods

To verify the generalizability of scRL’s superior fate decision intensity, we evaluated various machine learning and deep learning dimensionality reduction methods on two datasets as alternative approaches for fate decision intensity analysis, comparing them with the fate decision intensity obtained by scRL. In the hematopoietic and pancreatic datasets, we utilized the grid embeddings and pseudotime information obtained by scRL for subsequent fate decision intensity analysis (Fig. 4A-B). Using methods such as Diffusion Map, FA, ICA, LscVI, NMF, PCA, and scVI, we took the 95th, 90th, 85th, and 80th percentile intensities as alternative measures for fate decision evaluation and compared them with the fate decision intensity from scRL. Repeating experiments under sequences of highly variable genes ranging from 1000 to 5000, and evaluating with metrics like ARI, NMI, ASW, Davies-Bouldin, and Calinski-Harabasz, we found that scRL exhibited the most outstanding interpretability and achieved the highest overall score (Fig. 4C-D).

**Figure 4:**
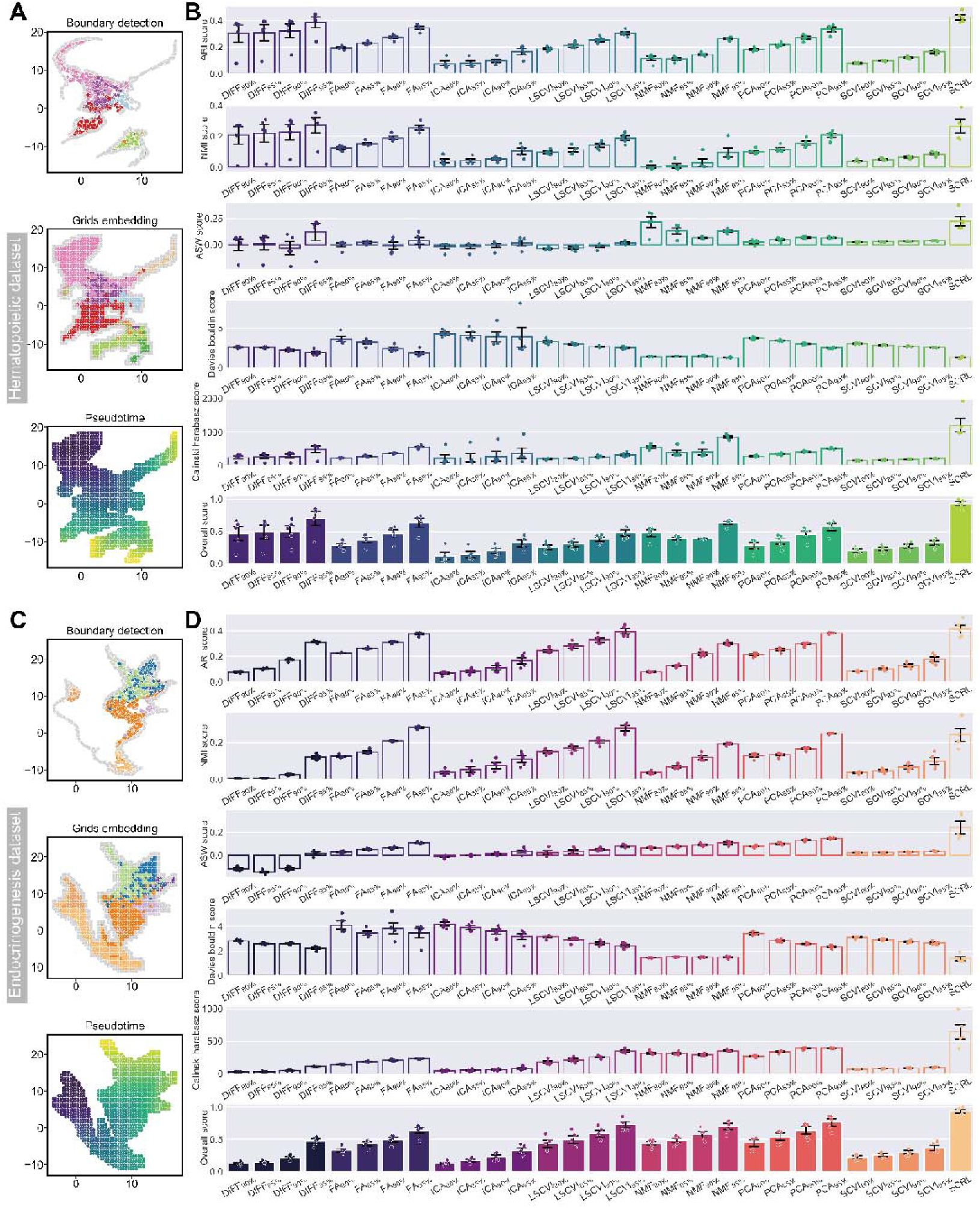
scRL Exhibits Superior Interpretability in Fate Decision Intensity Compared to Various Dimensionality Reduction Methods. (A, C) Boundary detection, grid embedding, and pseudotime inference in hematopoietic and endocrinogenesis datasets. (B, D) Comparison of interpretability metrics for scRL fate decision intensity against DIFF, FA, ICA, LscVI, NMF, PCA, and scVI at 95%, 90%, 85%, and 80% percentile intensities in hematopoietic and endocrinogenesis datasets.

### scRL Provides an Alternative Approach for Inferring Lineage Intensity

When lineage probabilities in single-cell data are unavailable or cannot be mapped to subgroup trajectory branches, scRL offers an alternative solution. This approach is also based on reinforcement learning, using labels from single-cell data (e.g., KMeans clustering labels) as reward signals to guide the agent in learning differentiation strategies, thereby evaluating the intensity of lineage differentiation. In the hematopoietic and pancreatic datasets, we employed KMeans clustering to obtain labels for five subgroups and used scRL to compute the lineage contribution intensity corresponding to each label. We observed that the lineage contribution intensity derived from scRL exhibited subgroup specificity within each label (Fig. 5A-B). Subsequently, to assess the interpretability of the contribution intensity obtained by scRL, we compared the latent spaces obtained by scRL with those from techniques such as PCA, ICA, FA, NMF, Diffusion Map, scVI, and LscVI. Under metrics like ASW, Davies-Bouldin, and Calinski-Harabasz, scRL demonstrated superior interpretability. Additionally, scRL’s lineage contribution intensity achieved the highest score in the overall assessment (Fig. 5C-D).

**Figure 5:**
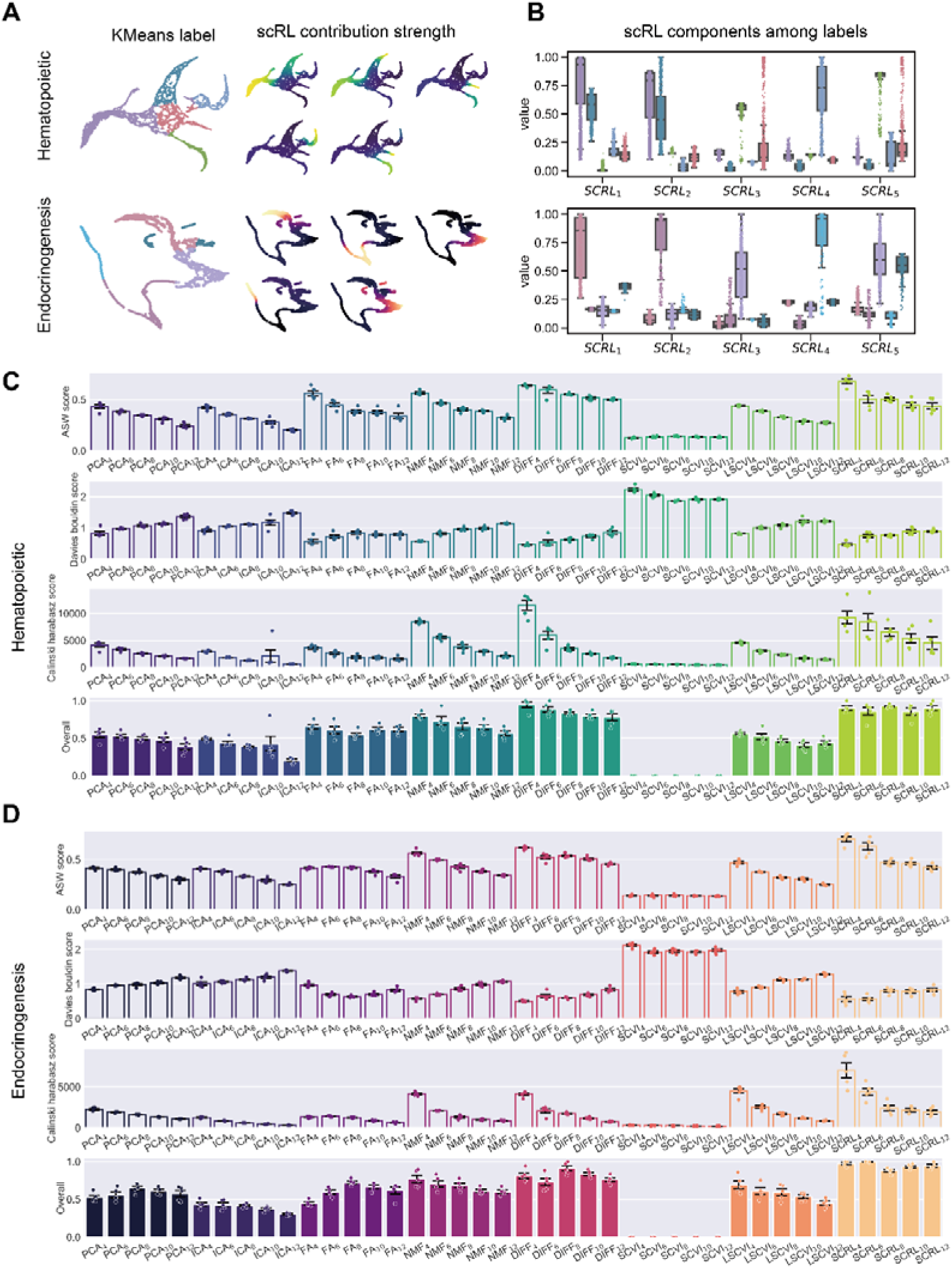
scRL Constructs Superior Lineage Intensity Using Subgroup Information Compared to Various Methods. (A) Distribution of 5 subgroups under KMeans and the distribution of subgroup lineage intensity obtained by scRL on UMAP. (B) Intensity of lineage components across different subgroups as determined by scRL. (C-D) Comparison of PCA, ICA, FA, NMF, Diff, scVI, LscVI, and scRL using ASW, Davies-Bouldin, and Calinski-Harabasz metrics. The comparison is conducted with KMeans subgroup numbers set at 4, 6, 8, 10, and 12, and highly variable gene sequences ranging from 1000 to 5000 as repetitions.

### The lineage and gene decision state value of scRL decode a common differentiation stage

To perform a detailed analysis of both gene and lineage decision state values, pseudotime analysis of scRL was conducted for a AML dataset^19^. We first identified four distinct branches representing lineage decision values (Fig. 6A-C). Then, we selected 10 marker genes as indicators of gene decision values for each of these branches, followed by gene decision analysis on these branch-specific genes and lineage decision analysis on four identified branches (Fig. 6D-F). Notably, a strong correlation between lineage and gene decision values was revealed within each branch (Fig. 6G). Additionally, it was also observed that the weighted average pseudotime of lineage decision values consistently preceded that of gene expression across all four branches (Fig. 6H).

**Figure 6:**
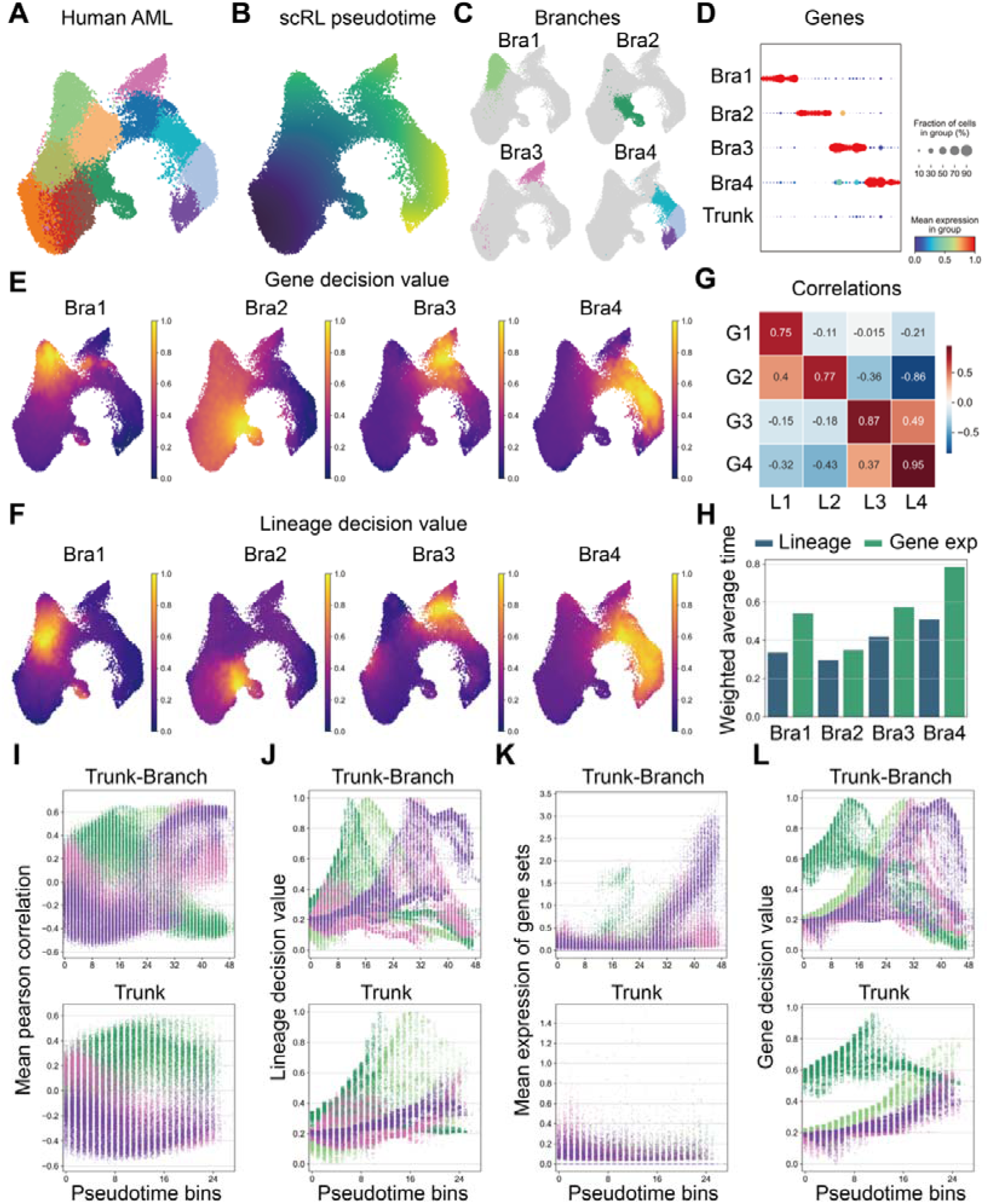
Combining lineage and gene information to decode fate decision states. (A) UMAP of human acute myeloid leukemia (AML), colored with clusters. (B) Pseudotime inferred by scRL. (C) 4 Branches of the UMAP space. (D) 10 marker genes sorted by log-foldchanges for each branch, with at least 25% percent cells express but less than 25% cells express in reference cell. (E) Gene set decision value of each branch. (F) Lineage decision value of each branch. (G) Pearson correlation heatmap of each branches’ gene set decision value represented by G1-4 and lineage decision value represented by L1-4. (H) Average pseudotime of the dataset weighted by the lineage decision value as well as the gene expression of each branch. (I) Mean Pearson correlation with branch specific cells along uniformly 50-binned pseudotime of the whole dataset at top panel and the trunk part at bottom panel. (J) Lineage decision value of each branch along uniformly 50-binned pseudotime of the whole dataset at top panel and the trunk part at bottom panel. (K) Mean expression of branch specific gene sets along uniformly 50-binned pseudotime of the whole dataset at top panel and the trunk part at bottom panel. (L) Gene decision value of branch specific gene sets along uniformly 50-binned pseudotime of the whole dataset at top panel and the trunk part at bottom panel.

Examination of the lineage decision values across the four branches, when mapped along binned pseudotime, revealed more distinct trajectory dynamics compared to the mean Pearson correlation values with cells specific to each branch (Fig. 6I-J). Similarly, the analysis of gene decision values along binned pseudotime provided a clearer depiction of the dynamics than the Pearson correlation values with branch-specific genes (Fig. 6K-L). This enhanced clarity in the data facilitated a deeper understanding of the developmental trajectories in the context of both lineage and gene decision processes. Our findings suggested that lineage decision values serve as early indicators of gene expression changes, providing valuable insights into the dynamics of gene regulation within the context of AML progression. And the approach underscored the utility of integrating pseudotime analysis with decision values to elucidate complex biological patterns and regulatory mechanisms in developmental processes.

## Discussion

The adaptation of reinforcement learning methodologies to biological systems highlights the versatility and potential of these techniques in addressing complex challenges in biological research.^23,24^In this study, our novel framework leverages the power of reinforcement learning to model and analyze cellular differentiation pathways within the high-dimensional landscape of single-cell data. By efficiently exploring and exploiting differentiation pathways, scRL identifies key nodes and pathways that influence cell fate decisions, providing valuable insights into cellular dynamics and potential avenues for manipulating cell fate in various biological applications.

Compared to traditional methods that rely on deterministic models or predefined genetic markers, such as CEFCON^25^, scRL offers a more dynamic and adaptive approach to analyzing cell fate decisions. The integration of reinforcement learning into single-cell data provides a nuanced analysis of differentiation pathways, including learning which pathways lead to optimal outcomes and identifying critical decision points within those pathways. This capability sets scRL apart from clustering algorithms or trajectory inference techniques that may not effectively pinpoint exact decision nodes^26^. The reward mechanism in scRL, based on lineage-specific and gene-specific targets, allows for continuous learning and adjustment of predictions. However, further improvements can be made to scRL’s reward modeling by integrating multi-modal data sources to enhance the granularity and accuracy of the reward signals.

In terms of trajectory inference, scRL’s module can be compared to other methods such as Monocle, TSCAN, and Slingshot^27^. Although these algorithms have their strengths, they may struggle with complex trajectory structures, miss transitional states, or be sensitive to noise and parameter settings. By integrating adaptive learning rates and incorporating feedback mechanisms based on experimental validations, we achieved the goal of improving the accuracy and robustness of scRL’s trajectory inference.

The gene trend module in scRL is preferable to algorithms like Wanderlust or Wishbone, which can also track the changes in gene expression but may be limited to specific types of trajectories or lineage progressions^28^. By adaptively selecting relevant genes based on their impact on trajectory decisions and integrating external datasets for validation, scRL can provide more nuanced insights and help understand the broader biological implications of these trends.

The intrinsic logic behind scRL’s critic network lies in its comprehensive evaluation, including integrating lineage or gene expression reward signals with two-dimensional trajectory information constrained by the grid-embedded environment. Moreover, the evaluation represents an expected reward estimation that decays with the gamma parameter, indicating the expected lineage or gene rewards associated with a particular state under a specific decay rate. The estimation enables the assessment of lineage decision and pre-gene expression states, which exhibit strong correlations within the same lineage, mutually reinforcing and revealing cellular fate decision. However, it should be acknowledged that scRL’s state value estimation relies on the recovery of differentiation trajectories by dimensionality reduction methods, which may lead to variations in lineage fate decision information due to the chosen technique. The definition of starting and target subpopulations of a lineage plays a pivotal role in the final estimation results, as lineage information is influenced by the nearest neighbor graph constructed from the latent space and dimensionality reduction methods like UMAP and DIFFMAP. In conclusion, this study presents a transformative tool in the field of computational biology, offering an approach to decoding the lineage and cell fate decisions based on single-cell sequencing.

## Methods

### Grids embedding function

The *Grids from Embedding* function (Fig. S21 Algorithm 1) processes a 2-D (two-dimensional) embedding space *x* to produce a grid-based representation. This function is parameterized to allow flexibility in grid generation, boundary detection, and computational efficiency through parallel processing.

### Grid and spine generation

Initially, the function invokes *Generate Grids* (Fig. S21 Algorithm 2), which computes grid points across the embedding space. This involves determining the extremal coordinates of the space (right, left, top, bottom) and using these to create a mesh grid of points. Additionally, the function constructs spines at the boundaries of the grid, which are essential for subsequent boundary detection processes.

### Boundary detection

Using parallel processing, the function calculates boundary grids by evaluating the arc distances from each spine point to all points in the embedding space (Fig. S21 Algorithm 4). This is achieved through the *Get Arc Dist* function (Fig. S21 Algorithm 3), which computes arc tangent values and distances, grouping them to identify the maximum distances that define the boundaries.

### Mask generation

Depending on the observer number *j*, the function either recalculates spines or reuses the existing ones to determine grids that should be masked out from analysis. This is done to focus the analysis on regions of interest, improving both the efficiency and relevance of the results (Fig. S21 Algorithm 5).

### Adjacency and connectivity

After mask generation, the function establishes adjacency relationships between the remaining grid points using the *Get Adjacent* function (Fig. S21 Algorithm 6). This step is crucial for understanding the connectivity and layout of the space, which aids in further analyses like clustering or region-based statistics.

### Boundary refinement

Finally, the function refines the detected boundaries using the *Check Houndary* function (Fig. S21 Algorithm 7). This ensures that all boundary grids are accurately identified, incorporating any adjacent grids that may have been initially overlooked.

### Annotation projecting function

We assign annotations to each specific grid point by utilizing the information from the nearest point in the original embedding.

### Pseudotime alignment function

As depicted in Algorithm of Figure S22, to generate a pseudotime ordering, we select grid points annotated as belonging to the starting subpopulation. From these, we randomly sample n points and calculate the average Dijkstra shortest path length from these n points to all remaining grid points, which serves as the pseudotime.

In cases where a matched grid world encompasses two or more components, the component with the highest number of grids is considered the primary one and utilize for initial pseudotime computation. For the remaining components, their starting points are determined as the points that have the minimum distance to any point in the primary component. The pseudotime values for these remaining points are then calculated based on the shortest path lengths from their respective starting points, using the pseudotime of the nearest point in the primary component as the starting value. This process is repeated for all components, ultimately yielding pseudotime values for the entire dataset.

### Value projection function

As depicted in Figure S23, the value projection function calculates the weights *w* as a normalized exponential function of the squared distances *D*^2^ from a point to its nearest *N* grids. The distances are scaled by the variance σ^2^, which modulates the influence of each grid based on its proximity. This results in a weighted average where closer grids have a higher impact on the outcome:

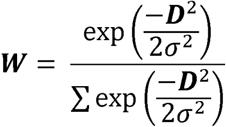

This function allows for smooth interpolation of values across the state space by calculating a weighted average of the nearest *N* grids, where the weights are determined by a Gaussian kernel based on the distance vector and standard deviation.

### Lineage reward function

For the lineage reward function, the target cluster grids are selected, and *t* represents the scaled pseudotime value. *β* is the decaying factor, *R_d_* is the decision mode reward, and *R_C_* is the contribution mode reward:

Decision Mode Reward:

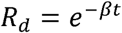

Contribution Mode Reward:

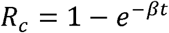

### Gene reward function

In the gene reward function, let *r* be the reward value, *P* be the penalty value, and *t* be the total pseudotime. *β* is the decaying factor. The decision mode reward *R_d_* and the contribution mode reward *R_C_* are calculated as:

Decision Mode Reward:

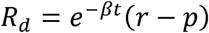

Contribution Mode Reward:

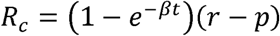

### Tabular Q learning

The Tabular Q-Learning algorithm is a form of reinforcement learning that updates the Q-values, which are estimates of the expected future rewards for a given state-action pair. The Q-values are stored on a table, with each entry corresponding to a state-action pair. The update rule for the Q-values is as follows:

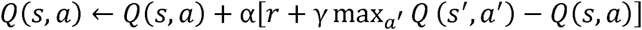

Here, α is the learning rate, *r* is the immediate reward, γ is the discount factor, and s’ is the next state. The update rule modifies the Q-value of the current state-action pair by incorporating the immediate reward plus the discounted maximum future reward, adjusted by the learning rate α. This ensures smooth convergence by scaling the difference between the updated and current Q-values.

### Actor-critic architecture

In the actor-critic architecture, the TD (temporal difference) error δ_t_ at time *t* is calculated as:

Where:

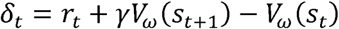

*r_t_* is the reward received after taking an action at state *s_t_*,

γ is the discount factor,

*V_ω_ (st)* is the value function parameterized by ω at state *s*.

This TD error measures the difference between the predicted value and the actual return received, guiding the updates for both the actor and the critic. The parameters ω and θ are updated as follows:

For the critic (updating ω):

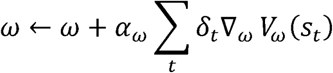

For the actor (updating *θ*):

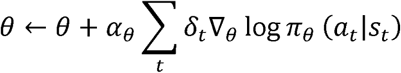

Here:

α_ω_ and α_θ_ are the learning rates for the critic and actor, respectively, 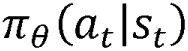 is the policy function parameterized by *θ*, 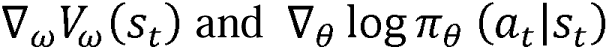 are the gradients with respect to their parameters.

### Autoencoder architecture for simulating

The encoder component of the autoencoder, denoted as *E*, processes inputs through individual neural network layers, which are then concatenated to form a unified latent representation *z*. This is mathematically represented as:

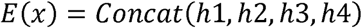

Where *x* includes *z_latent_,g_expr_,c_Coord_,t_pseudo_.* The decoder, D, then takes this concatenated latent space as input and aims to reconstruct the normal distribution of one-step differences for the encoder’s inputs, outputting:

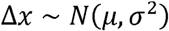

This approach allows the model to capture the dynamics of the input data by focusing on changes rather than absolute values. The primary objective of our training process is to minimize the reconstruction error of the one-step differences, thereby effectively capturing the dynamic changes in the input spaces. This setup is particularly advantageous for simulating and understanding complex biological processes, such as cellular development or changes in gene expression over time, where accurate modeling of stepwise trajectories is crucial.

### Advantages of scRL

As depicted in Figure S24 the functionalities of current pseudotime analysis tools could be broadly categorized into three areas: pseudotime and differentiation potential analysis, trajectory inference, and gene trends analysis. Most available packages estimated global pseudotime, with exceptions such as FateID^29^, which primarily focused on fate commitment value estimation, and Slingshot^11^, which emphasized pseudotime and trajectory inference based on principal curves.

While pseudotime alignment is not unique to scRL, our method demonstrates superior performance under specific conditions (Fig. S25C), positioning it as a competitive option in the rapidly evolving landscape of single cell pseudotime analysis tools.

In lineage potential analysis, only a few methods, such as Palantir, FateID, and scVelo, perform such estimations. Notably, scVelo requires both spliced and unspliced transcriptomic data, limiting its applicability. Palantir and FateID utilize supervised learning approaches, while scRL introduces a novel deep reinforcement learning-based method. Importantly, scRL’s lineage potential estimation strongly correlates with the embedding’s manifold (Fig. S25B), effectively leveraging trajectory information.

When assessing differentiation potential, scRL adopts a biologically meaningful approach by designating early-stage or stem cells as both reward lineages and starting points, simulating the fate decision process of stem cells. This contrasts with methods like Palantir and FateID, which rely on the entropy of already estimated lineage potentials.

Regarding trajectory inference, the Monocle series prefers the principal trees method, whereas Slingshot and FateID favor the principal curves approach. Both methods provide representations of the original data’s 2-D embedding space, showcasing the data generation process. However, by returning to the essence of the trajectory itself and employing the policy network to sample paths depicting the differentiation process, scRL introduces a fresh perspective to these techniques. A comparative analysis between our method and existing techniques has been conducted (Fig. S27). Innovatively, scRL can simulate the differentiation process beyond the original data scope.

For gene trend analysis, nearly all methods can perform comprehensive pseudotime alignment and predict gene trends through a general additive linear model (GAM). Similarly, scRL sets gene expression as the reward and utilizes the critic network to predict the gene decision state value, which can be defined as the intensity of the pre-gene expression state.

### Application of grid embedding and superiority of scRL pseudotime

Different embeddings offer distinct perspectives on the differentiation process. Here, we explore the versatility of our grid embedding function by applying it to six diverse dimensional reduction methods: PCA, diffusion map, UMAP, t-SNE, FA (force atlas), and Phate. Remarkably, each of these methods achieves impressive results when combined with grid embedding (Fig. S26A). Furthermore, the pseudotime alignment is seamlessly integrated with each grid embedding, providing a comprehensive view of the data (Fig. S26B).

To validate the effectiveness of scRL’s pseudotime, we conducted correlation analyses using Pearson, Spearman, and Kendall metrics. The results demonstrate that scRL’s pseudotime exhibits the highest correlation with the distance to the starting point on each embedding, which was established as ground truth (Figure S20C). And the superior could be interpreted as smoother pseudotime is estimated regarding each embedding.

### Comparison of lineage potential between methods

In lineage potential calculation, several methods have emerged, including FateID—a random forest-based supervised learning approach—and Palantir, which uses a diffusion map-based unsupervised method. scRL leverages deep reinforcement learning and a critic neural network to estimate potential. To assess the temporal association of lineage potential, we selected four well-distinguished hematopoietic branch embedding spaces: diffusion map, UMAP, FA, and PHATE (Fig. S26A).

Correlation analysis revealed that scRL’s lineage potential consistently achieves superior temporal correlation scores, indicating that the potential changes smoothly with time. In contrast, Palantir and FateID approaches are negatively associated with time, indicating a more approximate estimation (Fig. S26B). This comparison highlights scRL’s effectiveness in capturing temporal dynamics

### Distinctive Trajectory Representations Across Methods

scRL’s trajectory sampling is influenced by the gamma parameter, allowing for adjustable path shrinkage or divergence (Fig. S28, Fig. S29). Compared to other methods, scRL’s trajectory is based on grid embedding paths, showcasing all possible routes from source to target in a 2-D embedding.

In contrast, Monocle2 relies on a reverse graph embedding of the original data, resulting in a potentially unitary path. Monocle3’s path is derived from the UMAP embedding’s principal tree, which lacks diversity. Slingshot outlines the backbone of differentiation using principal curves of any 2-D embedding, resembling Monocle2’s approach. PAGA, a cluster-based method, presents a coarse-grained neighborhood structure among clusters.

While each method offers valuable insights, scRL’s adaptability to different embeddings and its ability to simulate authentic differentiation paths from start to maturity constitute its innovative advantage as scRL is designed for optimizing a policy neural network and sampling a real trajectory on any embedding.

### Parameters

*n* (Grid Density Parameter):

The parameter *n* represents the density of the background grid to be generated in the preprocessing step of scRL. Specifically, it generates an *n* x *n* grid embedding based on the circumscribed rectangle of the input two-dimensional embedding of single cells.

*j* (Observer Number):

The parameter *j* denotes the number of observation points for grid embedding in the preprocessing step of scRL. The grid embedding function generates *j* observation points equidistantly on the four boundaries of the circumscribed rectangle of the original 2-D data embedding to scan the edges of the original data. More observation points indicate finer edge detection.

*γ* (Discount Factor):

The parameter *γ* is a factor in the reinforcement learning training process of scRL, representing the discounting degree of future rewards during the policy iteration process. Its value ranges from 0 to 1. A higher value indicates greater emphasis on future rewards, making the agent more forward-looking. However, too low a value may prevent the agent from learning the optimal differentiation strategy, resulting in significant deviations in the evaluation of state value.

α (Learning Rate):

The parameter α represents the learning rate during the policy iteration process, indicating the acceptance of new observations. The default learning rate for the policy network optimizer is set to 0.005, while the default learning rate for the critic network optimizer is set to 0.0001.

*n_neighbor_* (Neighbor Number):

The parameter *n_neighbor_* represents the number of neighbors considered when mapping between the grid embedding and the original embedding.

### Parameters robustness

We have conducted experiments to examine the robustness of scRL’s key parameters. Primarily, we focused on parameters *n* and *j* within the grid embedding function. Here, *n* represents the width of the anticipated grid world; a higher value of *n* indicates a denser grid environment. Meanwhile, *j* can be interpreted as the number of observers surrounding the grid. As the number of observers increases, the generated boundary becomes more precise (Fig. S30, Fig. S31).

It is important to note that excessively high *j* value could potentially introduce inconsistencies within the grid world, which may make certain island areas inaccessible to the agent, thereby rendering their state values unavailable. To assess the stability of pseudotime estimations, we tested various values of parameter *n* for each value of the parameter *j* and found that the results remained stable across different groups of parameters *j* (Figure S32A and S32B).

### Calculation of correlation coefficient

For the pseudotime correlation with the distance-based ground truth, we calculated the Pearson, Spearman, and Kendall correlation coefficients using the Pandas package after compiling the pseudotime and distance into one data frame. For the temporal correlation of gene expression values or lineage contribution values, the Pearson correlation coefficient was calculated.

### Calculation of distance-based ground truth

The distance-based ground truth for cells is the Euclidean distance from a pre-defined starting cell in a certain 2-D embedding space.

### Calculation of weighted average time

The pseudotime is weighted by the expression value of genes or lineage decision value which the sum of weight contribution is 1.

### Calculation of metrics

The calculations for various evaluation metrics, such as NMI, ARI, ASW, Calinski-Harabasz index, Davies-Bouldin index, and Pearson correlation, are implemented using Python’s scikit-learn library.

### Repetition of evaluation experiments

Most machine learning methods do not follow the gradient descent-based optimization process. Therefore, in evaluations involving machine learning techniques, we conducted repeated experiments using highly variable genes obtained from log-normalized single-cell data. The sequence of high variable gene counts was set at 1000, 2000, 3000, 4000, and 5000.

## Data and code availability

The single-cell RNA-seq data have been deposited in the GEO database under accession numbers GSE277292 and GSE278673 and are publicly available as of the publication date. All original codes have been deposited at Github at https://github.com/PeterPonyu/scRL and is publicly available as of the date of publication.

## Acknowledgments

We gratefully acknowledge the support from the National Natural Science Foundation of China through grants 82430103, 81930090, and 81725019 (awarded to J.W.), as well as 82073487 and 81602790 (awarded to S.C.). Additionally, this research was funded by the High-level Talent Program of the Third Military Medical University under grant 2022XRC02 (awarded to S.C.).

## Author Contributions

Z.F. was instrumental in developing the software. K.S., Z.F., and S.W. carried out most of the experiments with skill and dedication, while B.X., X.Y., Z.W., M.S., M.C., and F.C. provided essential assistance with various experimental tasks. Y.X., S.C., and J.W. made significant contributions through insightful scientific discussions and were actively involved in the preparation of the manuscript.

## Declaration of interests

The authors declare no competing interests.

## Notes

https://github.com/PeterPonyu

https://www.ncbi.nlm.nih.gov/geo/query/acc.cgi?acc=GSE277292

https://www.ncbi.nlm.nih.gov/geo/query/acc.cgi?acc=GSE278673

### Competing Interest Statement

The authors have declared no competing interest.

### Summary of Updates

This revision includes extensive methodological performance comparisons, and the procedural description of scRL can be found in the supplementary file.

