## Supplementary figures for "scRL: Utilizing Reinforcement Learning to Evaluate Fate Decisions in Single-Cell Data"

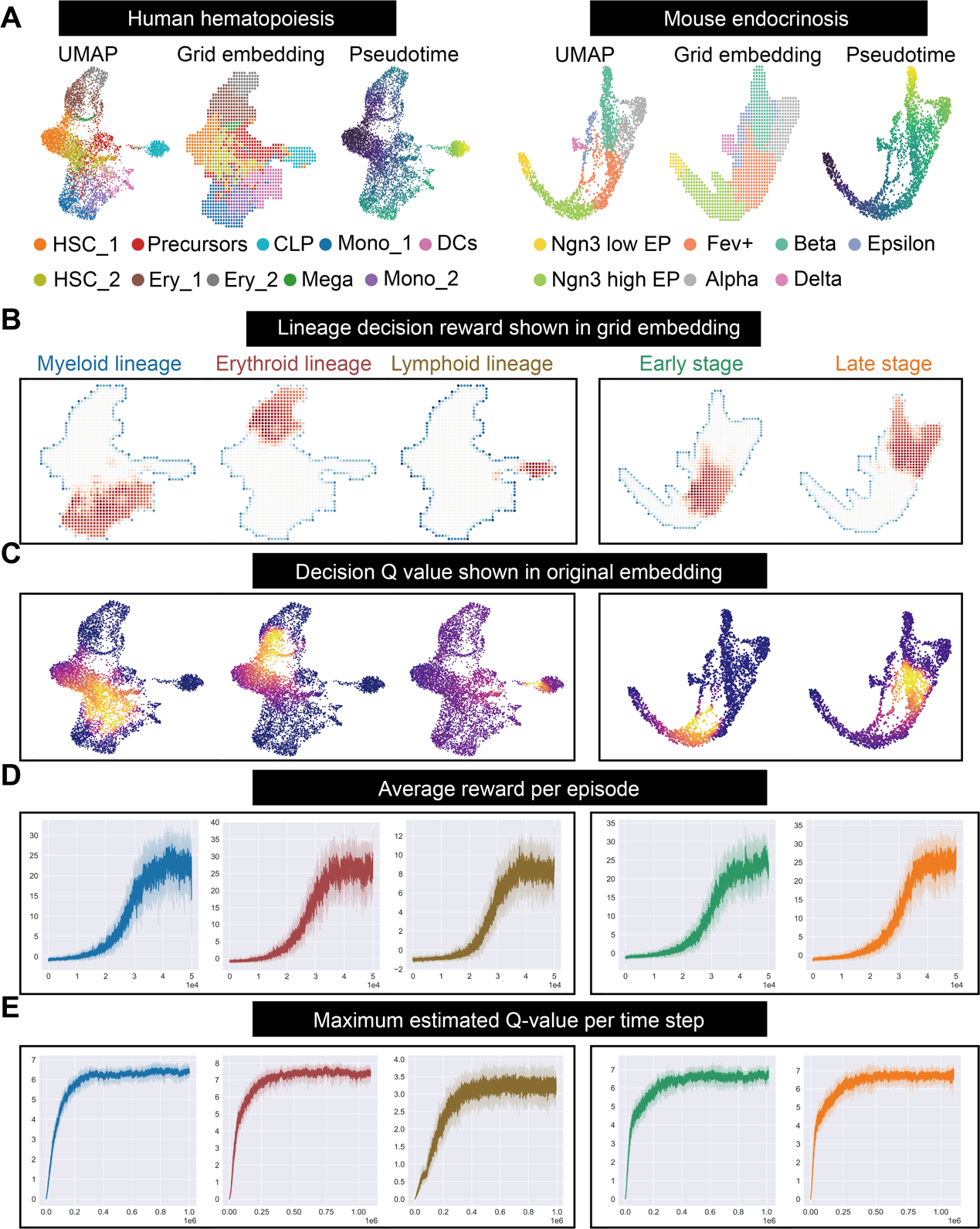
**Figure S1: The application of tabular Q learning on lineage decision reward mode grid world generated by scRL.**

(A) Preprocessing steps of grid embedding and pseudotime alignment applied to human hematopoiesis and mouse endocrinogenesis datasets enabling the representation of complex biological processes in a simplified grid world format. (B) Lineage decision rewards generated for each hematopoietic lineage and the various differentiation stages of mouse endocrinogenesis. (C) State values estimated by tabular Q-learning for each hematopoietic lineage and the differentiation stages of mouse endocrinogenesis. (D) The average reward per episode during the lineage decision reward mode training process. (E) The maximum state value estimated per updating step.

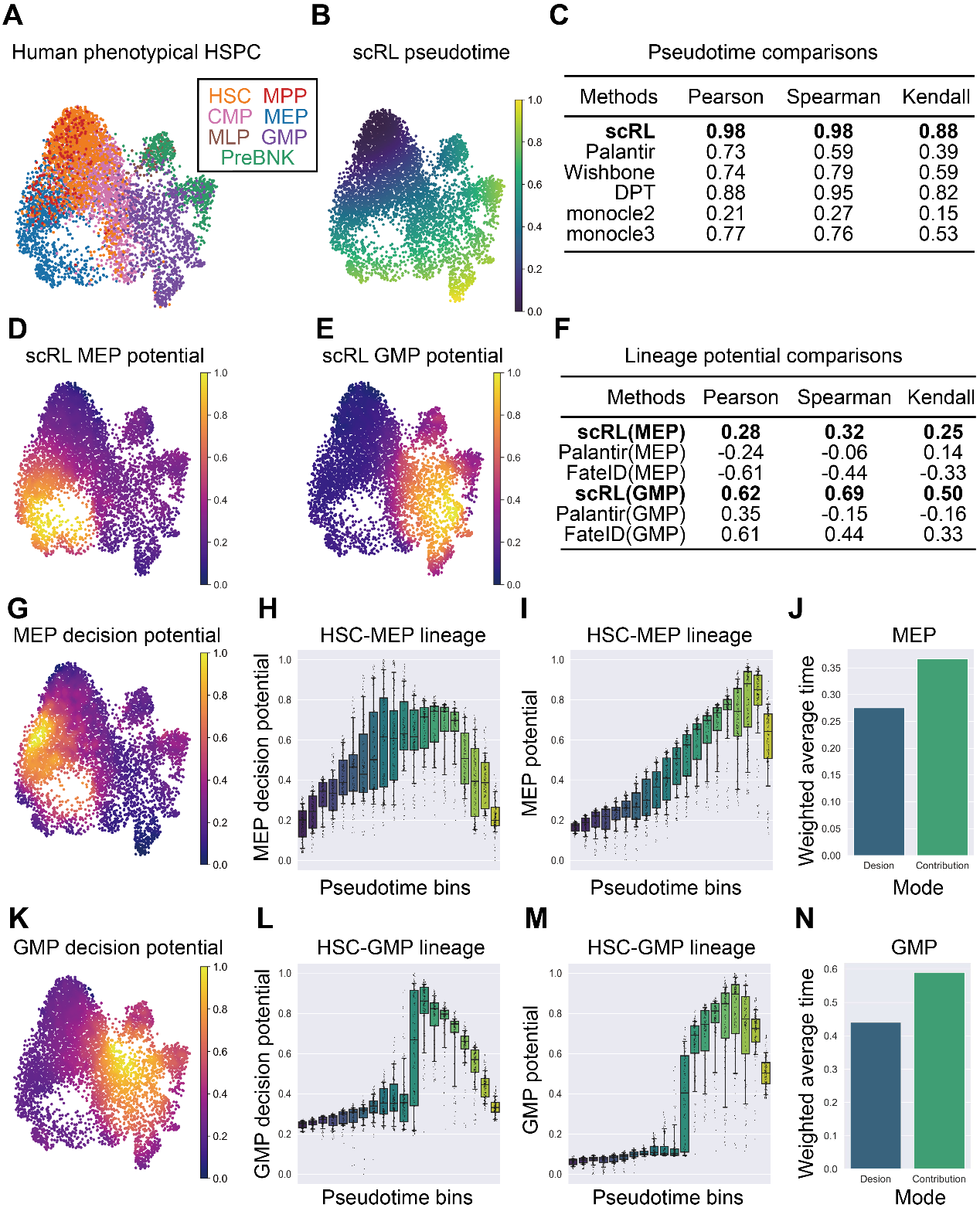

**Figure S2: Pseudotime and lineage potential function of scRL**

(A) UMAP projection of phenotypes of HSPC (hematopoietic stem and progenitor cell), including HSC (hematopoietic stem cell), MPP (multipotent progenitor), CMP (common myeloid progenitor), MLP (multipotent lymphoid progenitor), MEP (Megakaryocyte erythrocyte progenitor), GMP (Granulocyte monocyte progenitor), and PreBNK (Pre-B cell and Pre-natural killer cell). (B) Pseudotime aligned by scRL. (C) Comparisons between scRL and other methods including Palantir, Wishbone, DPT (diffusion pseudotime), Monocle2, Monocle3 with highest score in bold format. Pearson, Spearman and Kendall correlations are computed between pseudotime and the distance-based ground truth. The ground truth is defined as Euclidean distance from starting point in UMAP space. (D) MEP potential inferred by scRL with MEP cluster as reward and HSC cluster as start. (E) GMP potential inferred by scRL with GMP cluster as reward and HSC cluster as start. (F) Comparisons between scRL and other methods including Palantir and FateID with highest score in bold format. Pearson, Spearman and Kendall correlations are computed between MEP as well as GMP potential and the distance-based ground truth. The ground truth is defined as Euclidean distance from starting point in UMAP space. (G) MEP decision potential inferred by scRL with MEP cluster as reward and HSC cluster as start. (H-I) MEP potential and decision potential within HSC-MEP lineage along binned pseudotime. (J) Weighted average of pseudotime by MEP decision potential and lineage(contribution) potential. (K) GMP decision potential inferred by scRL with GMP cluster as reward and HSC cluster as start. (L-M) GMP potential and decision potential within HSC-GMP lineage along binned pseudotime. (N) Weighted average of pseudotime by GMP decision potential and lineage(contribution) potential.

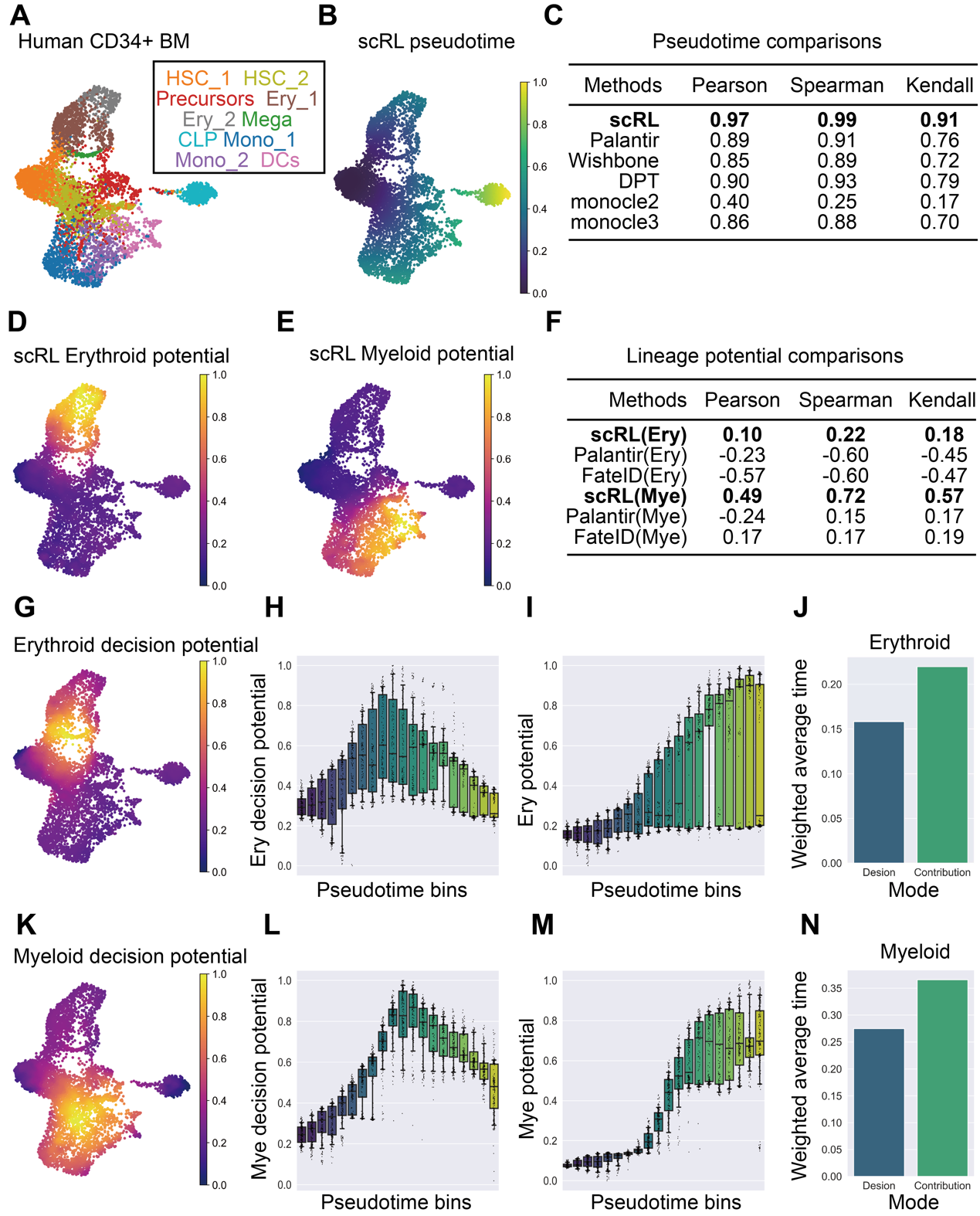

**Figure S3: Pseudotime and lineage potential function applied on human CD34^+^ bone marrow cells**

(A) UMAP projection of human CD34^+^ bone marrow cells, (B) Pseudotime aligned by scRL. (C) Comparisons between scRL and other methods including Palantir, Wishbone, DPT (diffusion pseudotime), monocle2, monocle3. Pearson, Spearman and Kendall correlations are computed between pseudotime and the distance-based ground truth. The ground truth is defined as Euclidean distance from starting point in UMAP space. (D) Erythroid potential inferred by scRL with Ery_1, Ery_2, Mega clusters as reward and HSC_1, HSC_2 cluster as start. (E) Myeloid potential inferred by scRL with Mono_1, Mono_2, DCs cluster as reward and HSC_1, HSC_2 cluster as start. (F) Comparisons between scRL and other methods including Palantir and FateID with highest score in bold format. Pearson, Spearman and Kendall correlations are computed between erythroid as well as myeloid potential and the distance-based ground truth. The ground truth is defined as Euclidean distance from starting point in UMAP space. (G) Erythroid decision potential inferred by scRL with Ery_1, Ery_2, Mega clusters as reward and HSC_1, HSC_2 cluster as start. (H-I) Erythroid potential and decision potential within erythroid lineage along binned pseudotime. (J) Weighted average of pseudotime by erythroid decision potential and lineage(contribution) potential. (K) Myeloid decision potential inferred by scRL with Mono_1, Mono_2, DCs cluster as reward and HSC_1, HSC_2 cluster as start. (L-M) Myeloid potential and decision potential within myeloid lineage along binned pseudotime. (N) Weighted average of pseudotime by myeloid decision potential and lineage(contribution) potential.

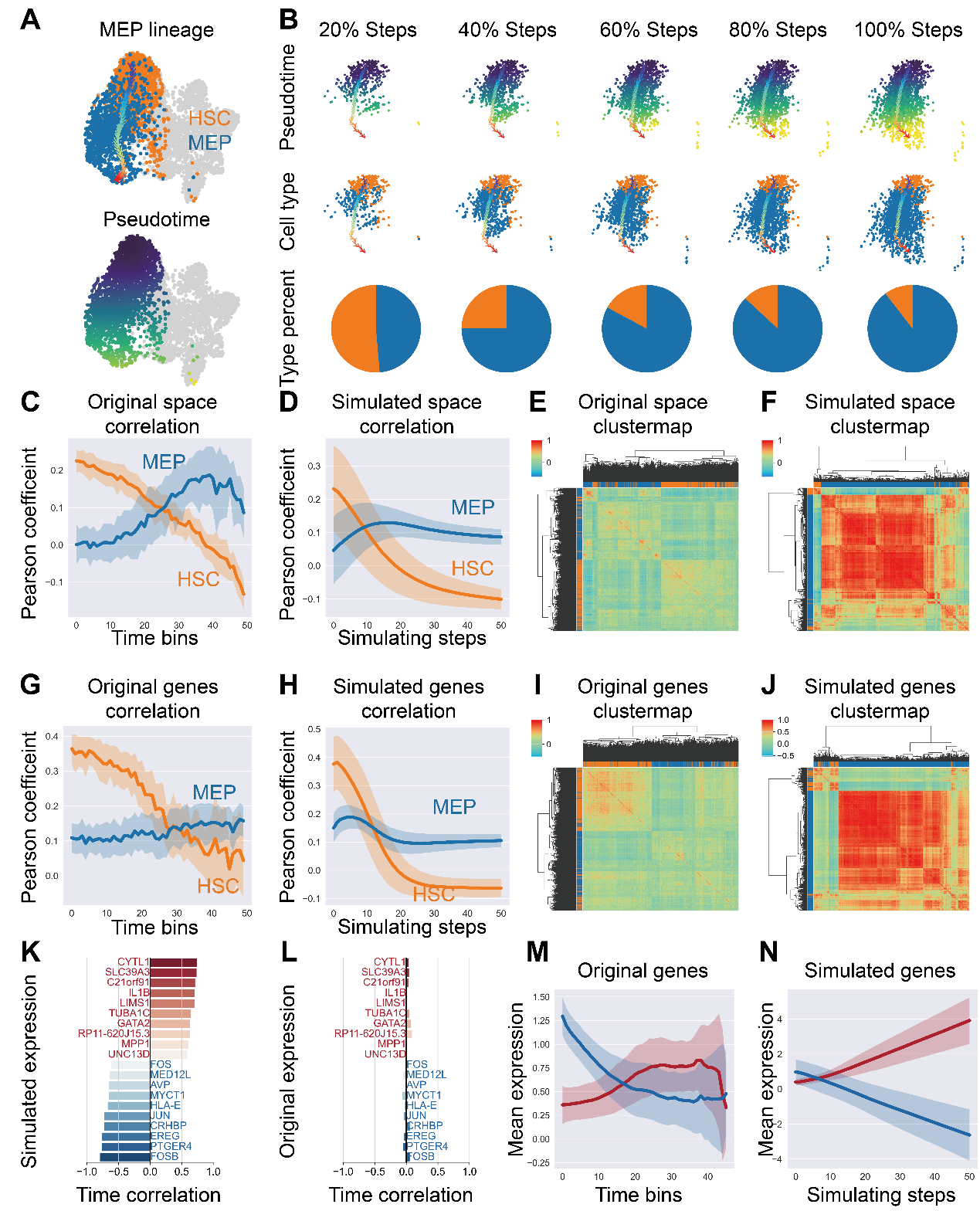

**Figure S4: Trajectory simulating function of scRL**

(A) MEP lineage and pseudotime aligned by scRL. (B) Simulated MEP trajectory with 20%, 40%, 60%, 80% steps and the whole trajectory. A single trajectory steps 50 times and 100 trajectories in total. Cell types are colored as shown in (A) and black dots referred to the starting points. Pie plots show the dynamic changes of cell type components. (C-D) Pearson correlation coefficient of the original and simulated PCA space of cells along uniformly 50-binned pseudotime, the mean value is represented by a line and the standard error is indicated by the colored filled area, examining the relationship between all cells in relation to MEP and HSC. (E-F) Hierarchical clustering of Pearson correlation between all the cells in original and simulated PCA space, cells are colored as in (A). (G-H) Pearson correlation coefficient of the original and simulated gene expression space of cells along uniformly 50-binned pseudotime. (I-J) Hierarchical clustering of Pearson correlation between all the cells in original and simulated gene expression space. (K-L) The correlation score with pseudotime of 10 most and 10 least relevant genes in simulated and original expression space. (M-N) The moving average of original and simulated mean expression along binned pseudotime of 10 most time relevant genes are colored as MEP, and 10 least relevant genes are colored as HSC, with line represents the mean value and filled area indicates the standard error.

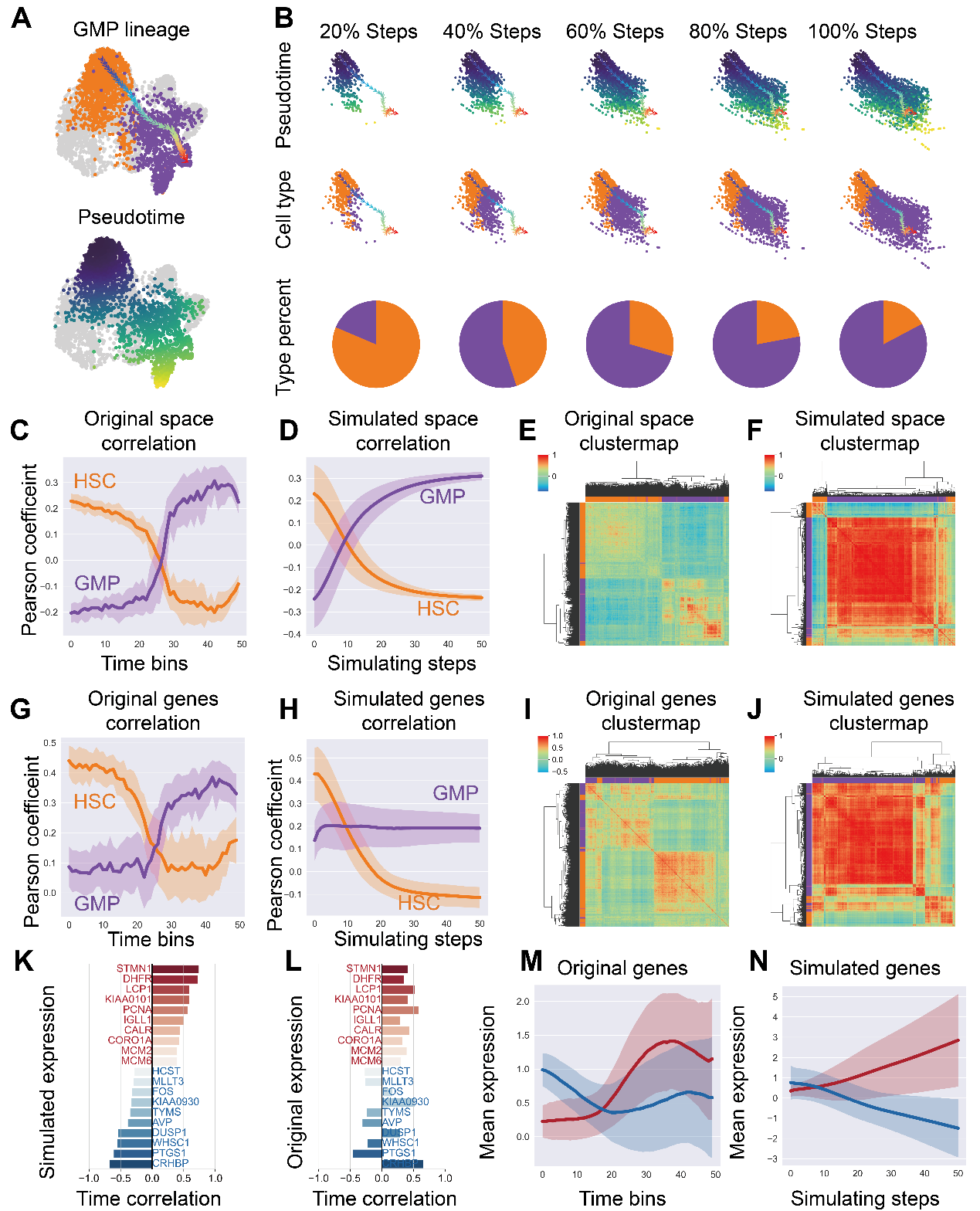

**Figure S5: Trajectory simulating function applied on phenotypic GMP lineage**

(A) GMP lineage and pseudotime aligned by scRL. (B) Simulated GMP trajectory with 20%, 40%, 60%, 80% steps and the whole trajectory. A single trajectory steps 50 times and 100 trajectories in total. Pie plots show the dynamic changes of cell type components. (C-D) Pearson correlation coefficient of the original space and simulated space of cells along uniformly 50-binned pseudotime, the mean value is represented by a line and the standard error is indicated by the colored filled area, examining the relationship between all cells in relation to GMP and HSC. (E-F) Hierarchical clustering of Pearson correlation between all the cells in original space and simulated space. (G-H) The Pearson coefficient along binned pseudotime and simulated steps calculated in gene expression space and simulated expression space. (I-J) Hierarchical clustering of Pearson correlation between all the cells in gene expression space and simulated expression space. (K-L) The correlation score with pseudotime of 10 most and 10 least relevant genes in simulated space and gene expression space. (M-N) The moving average of original mean expression and simulated mean expression along binned pseudotime of 10 most time relevant genes are colored as red, and 10 least relevant genes are colored as blue, with line representing the mean value and filled area indicates the standard error.

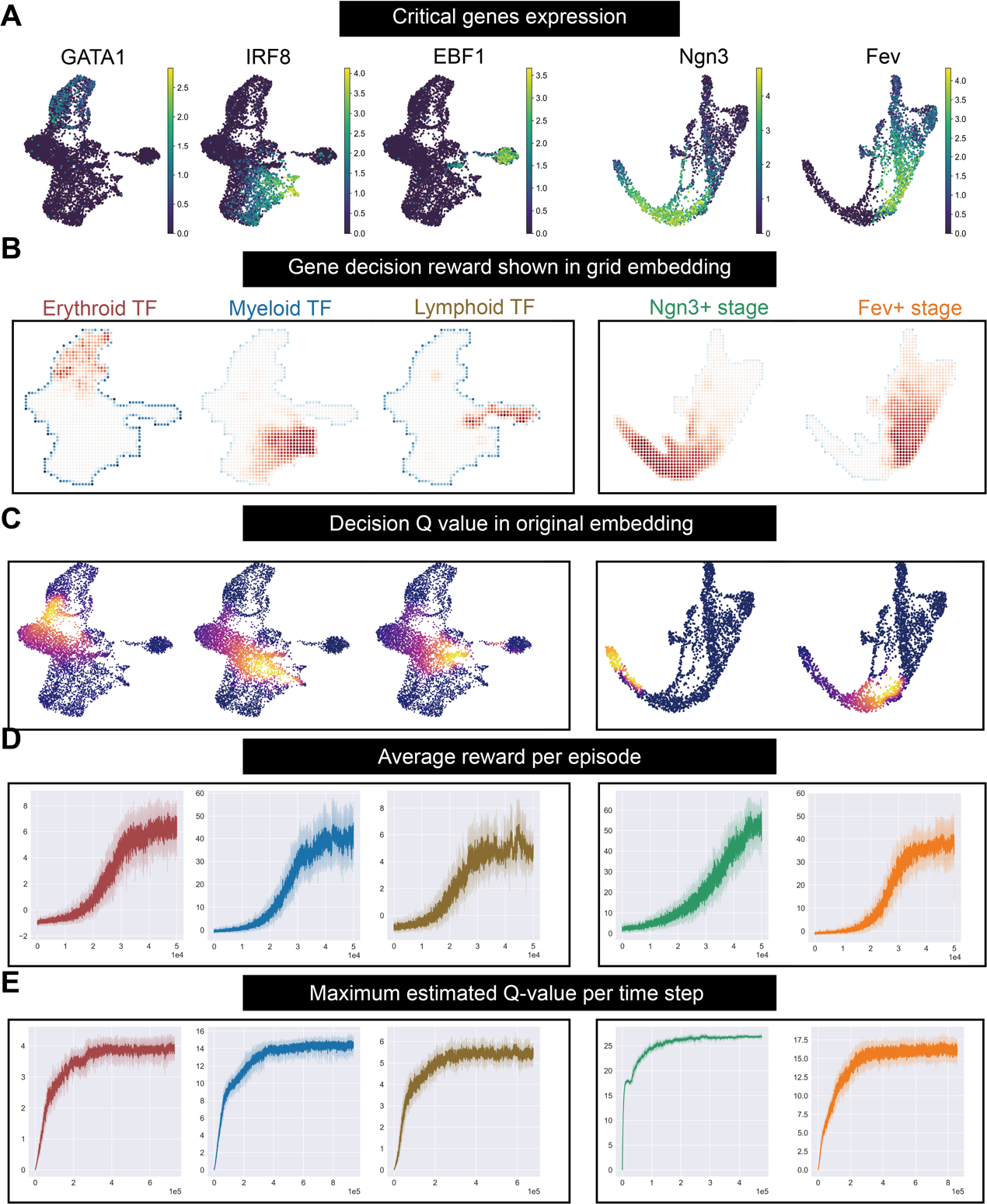

**Figure S6: The application of tabular Q learning on gene decision reward mode grid world generated by scRL.**

(A) Expression levels of critical transcription factors for each hematopoietic lineage and stage markers for endocrinogenesis. (B) Gene decision rewards generated for the genes. (C) State values estimated by tabular Q-learning for each gene, referring to the pre gene expression state intensity. (D) The average reward per episode during the gene decision reward mode training process. (E) The maximum state value estimated per updating steps.

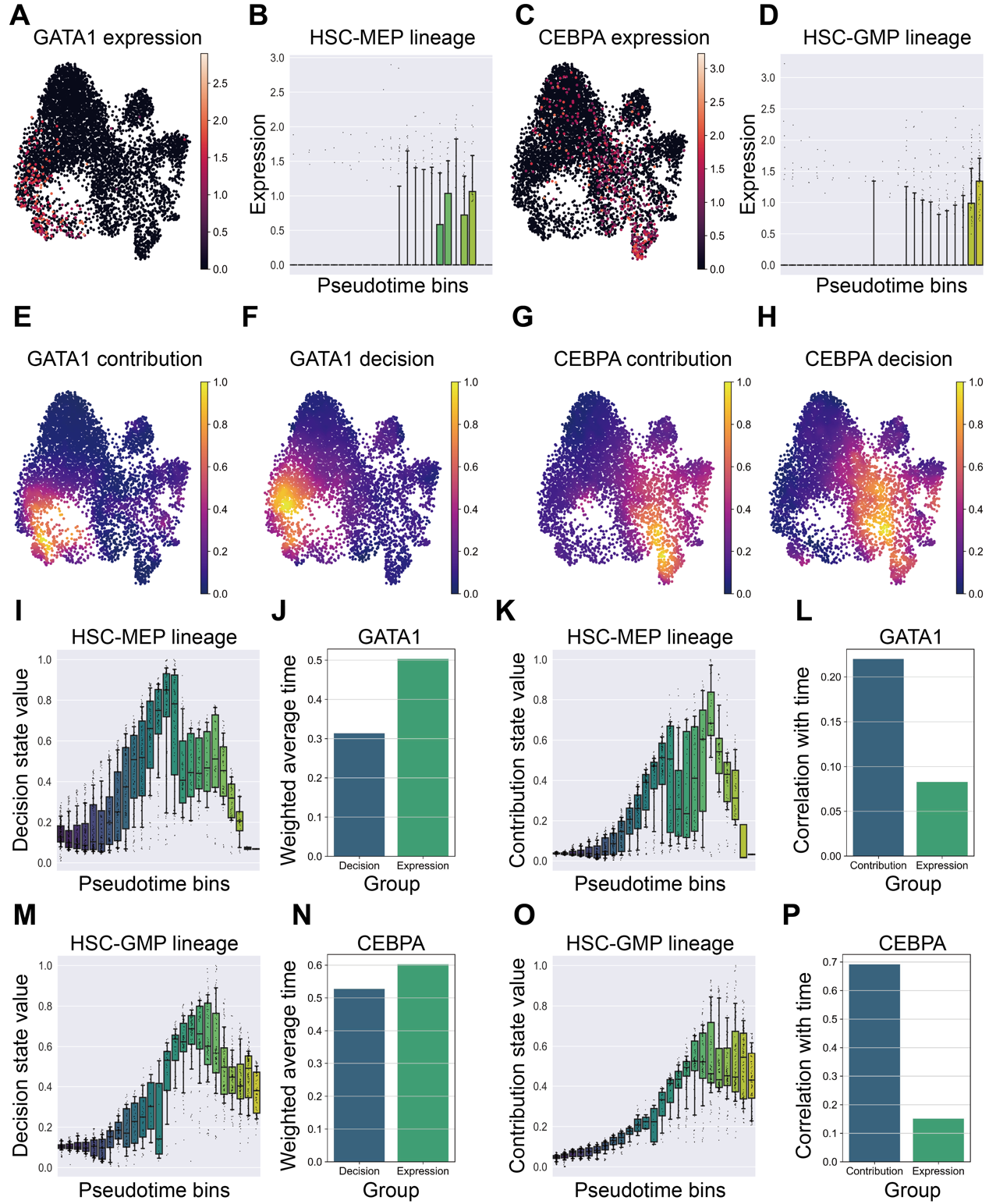

**Figure S7: Gene decision function of scRL**

(A) GATA1 expression on UMAP marking the erythroid lineage. (B) GATA1 expression on HSC-MEP (erythroid)lineage along binned pseudotime. (C) CEBPA expression on UMAP marking the myeloid lineage. (D) CEBPA expression on HSC-GMP (myeloid)lineage along binned pseudotime. (E) GATA1 gene contribution value on UMAP. (F) GATA1 gene decision value on UMAP (G) CEBPA gene contribution value on UMAP. (H) CEBPA gene decision value on UMAP. (I) The HSC-MEP lineage decision value along binned pseudotime. (J) The average pseudotime in HSC-MEP lineage weighted by GATA1 decision value and expression. (K) The HSC-MEP lineage contribution value along binned pseudotime. (L) The average correlation with pseudotime of GATA1 contribution value and its expression. (M) The HSC-GMP lineage decision value along binned pseudotime. (N) The average pseudotime in HSC-GMP lineage weighted by CEBPA decision value and expression. (O) The HSC-GMP lineage contribution value along binned pseudotime. (P) The average correlation with pseudotime of CEBPA contribution value and its expression.

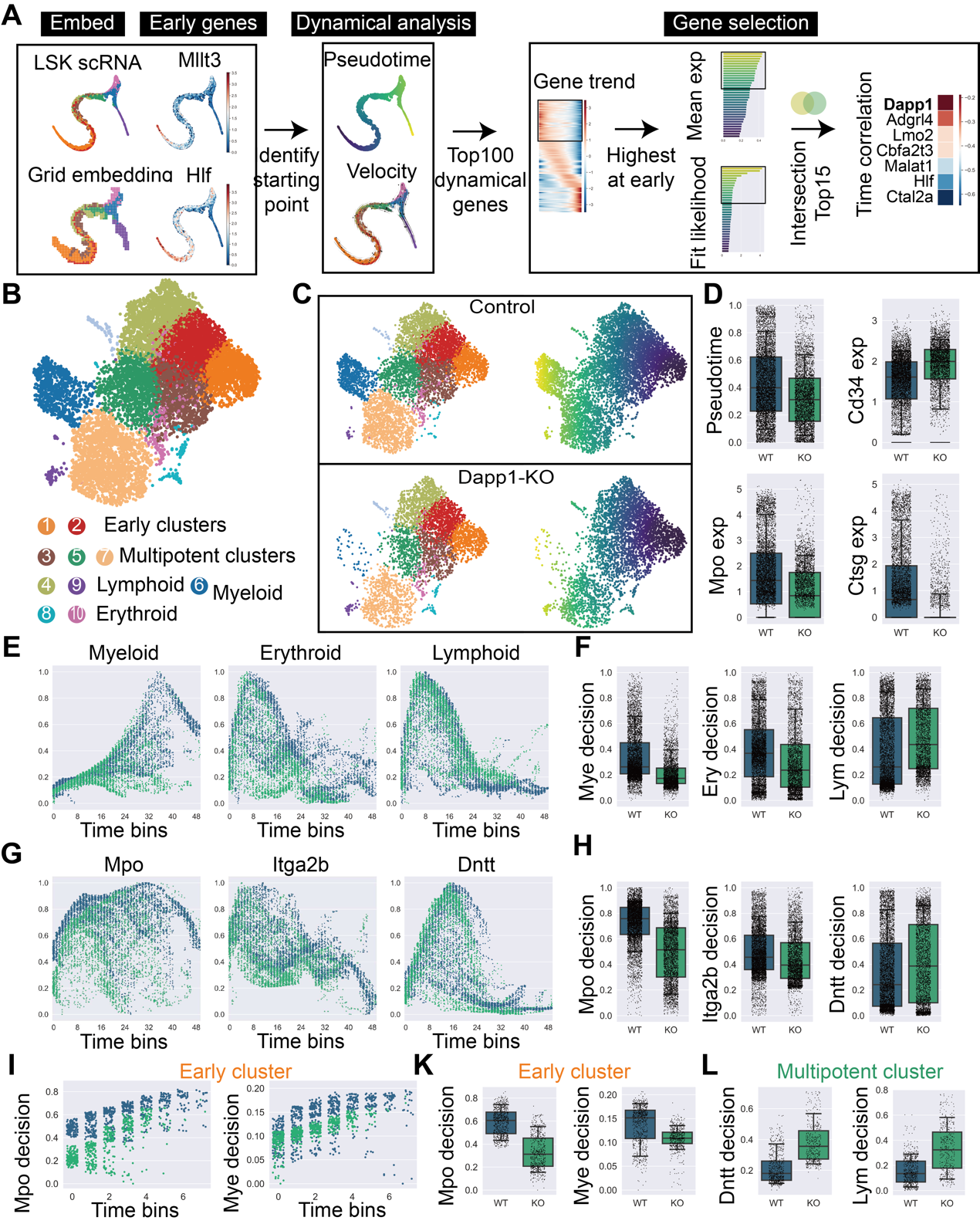

**Figure S8: Lineage commitment of HSCs at the earliest primitive stage perturbed by knockout of critical dynamical gene.**

(A) The workflow for identifying critical dynamical gene Dapp1 in the differentiation of HSCs begins from left panel with embeddings of LSK single-cell data to pinpoint the differentiation starting point. It then employs velocity analysis to identify the top dynamical genes involved in differentiation, meanwhile pseudotime is used to select the earliest expressing genes, from which the highest expressed and most dynamic genes are intersected. Finally, the right panel displays a time correlation score to rank the intersected genes. (B) Dapp1 knockout LSK atlas with clusters 1,2 annotated as Early clusters, 3,5,7 as multipotent clusters, 4, 9 as lymphoid clusters, 6 as myeloid cluster and 8, 10 as erythroid clusters. (C) Cluster distributions and pseudotime for control group and knockout group. (D) Box plots for comparisons of pseudotime and the expression of Cd34, Mpo, Ctsg. (E) Lineage decision value of myeloid, erythroid and lymphoid along uniformly 50-binned pseudotime. (F) Box plots for comparison of lineage decision value of myeloid, erythroid and lymphoid lineages. (G) Gene decision value of Mpo, Itga2b and Dntt along uniformly 50-binned pseudotime. (H) Box plots for comparison of gene decision value of Mpo, Itga2b and Dntt. (I) The myeloid lineage, Mpo decision value within early cluster 1 along binned pseudotime. (J) Box plots for comparison of myeloid lineage and Mpo decision value within early cluster 1. (K) Box plots for comparison of lymphoid lineage and Dntt decision value within multipotent cluster 5.

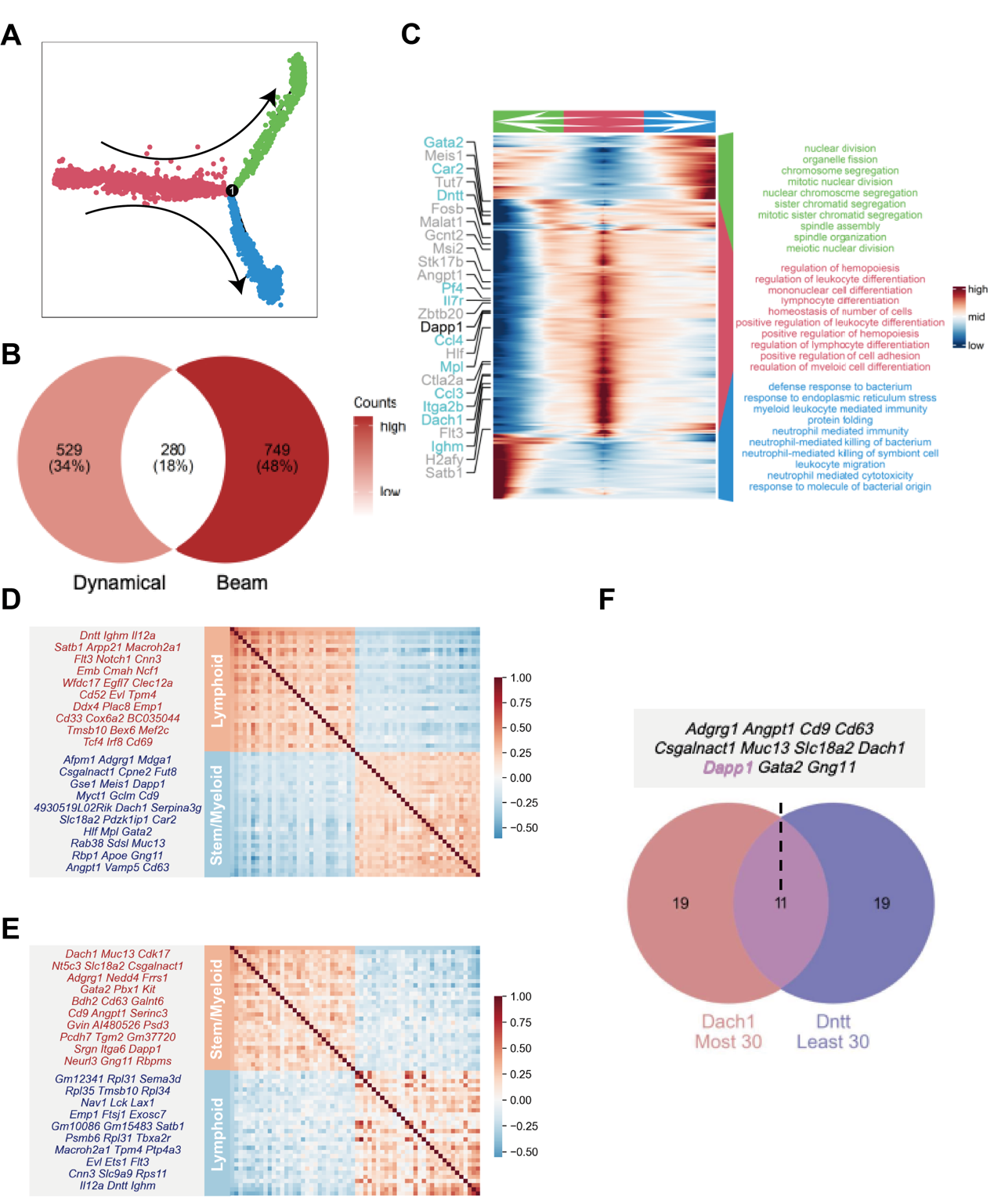

**Figure S9: Dapp1 serves as a differentiation dynamical gene in normal hematopoiesis**

(A) Monocle2 differentiation trajectory (B) Intersection of Dynamical genes and monocle2’s BEAM test significant genes, finally 280 genes were get where Dapp1 serve as an early gene. (C) Gene expression trend on different trajectories with GOBP terms enriched. (D) Gene correlation analysis of LSK single cell data from GSE136341. Dach1 most 30 correlated genes and least correlated genes were presented beside the heatmap. (E) Gene correlation analysis of LSK single cell data from GSE145491. Dntt most 30 correlated genes and least 30 correlated genes were presented beside the heatmap. (F) The intersection of Dach1 most correlated 30 genes and Dntt least correlated genes were presented.

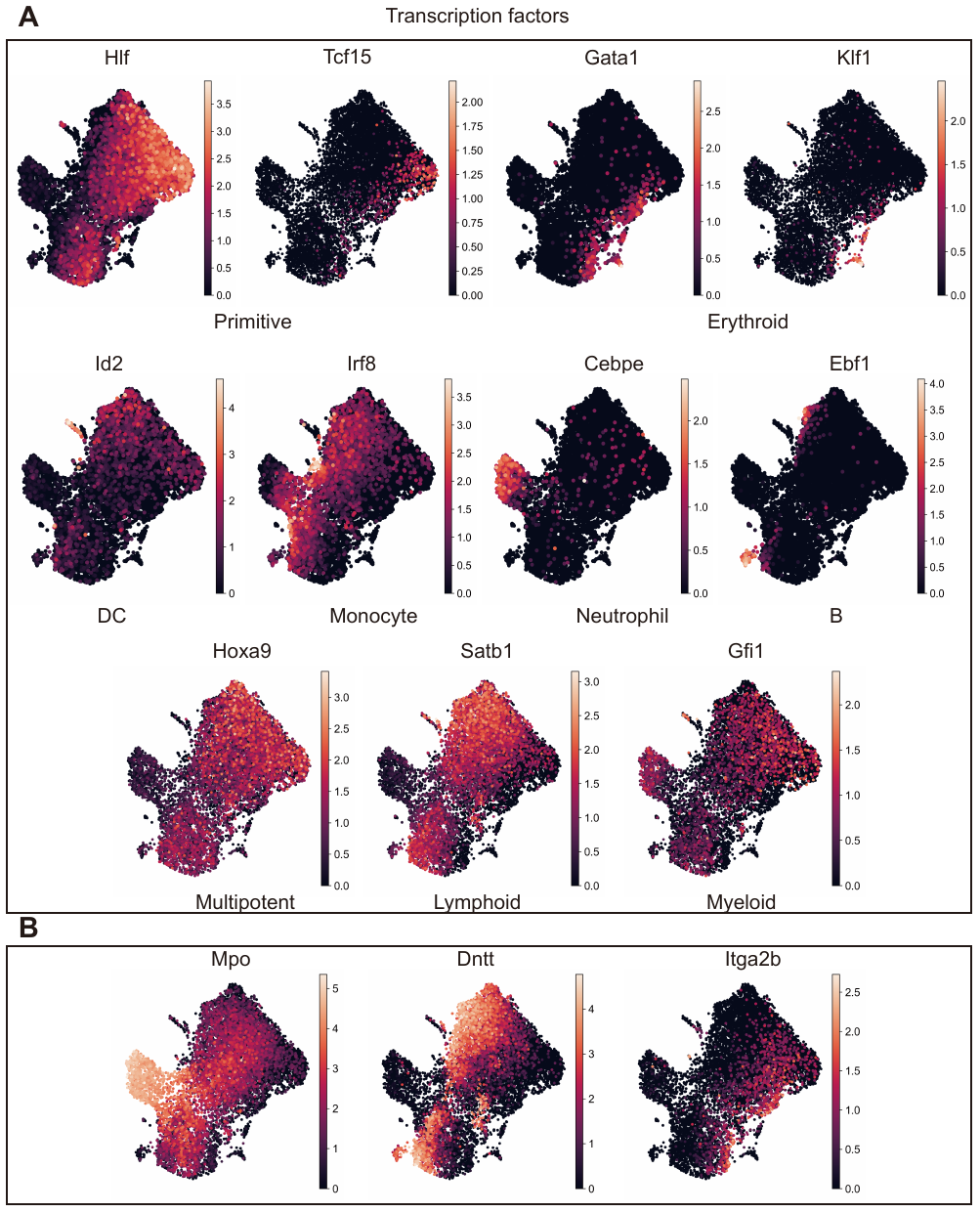

**Figure S10: Critical Gene Expression in Dapp1-Perturbed Datasets**

(A) Transcription factors’ expression critical for the identification of distinct hematopoietic lineages are presented on UMAP, including Hlf and Tcf15 for primitive HSCs, Gata1 and Klf1 for erythroid bias, Id2 for dendritic cells, Irf8 for monocytes, Cebpe for neutrophil, Ebf1 for B cells. Hoxa9 for multipotency, Satb1 for lymphoid bias and Gfi1 for myeloid bias. (B) The expression of Mpo, Dntt and Itga2b on UMAP.

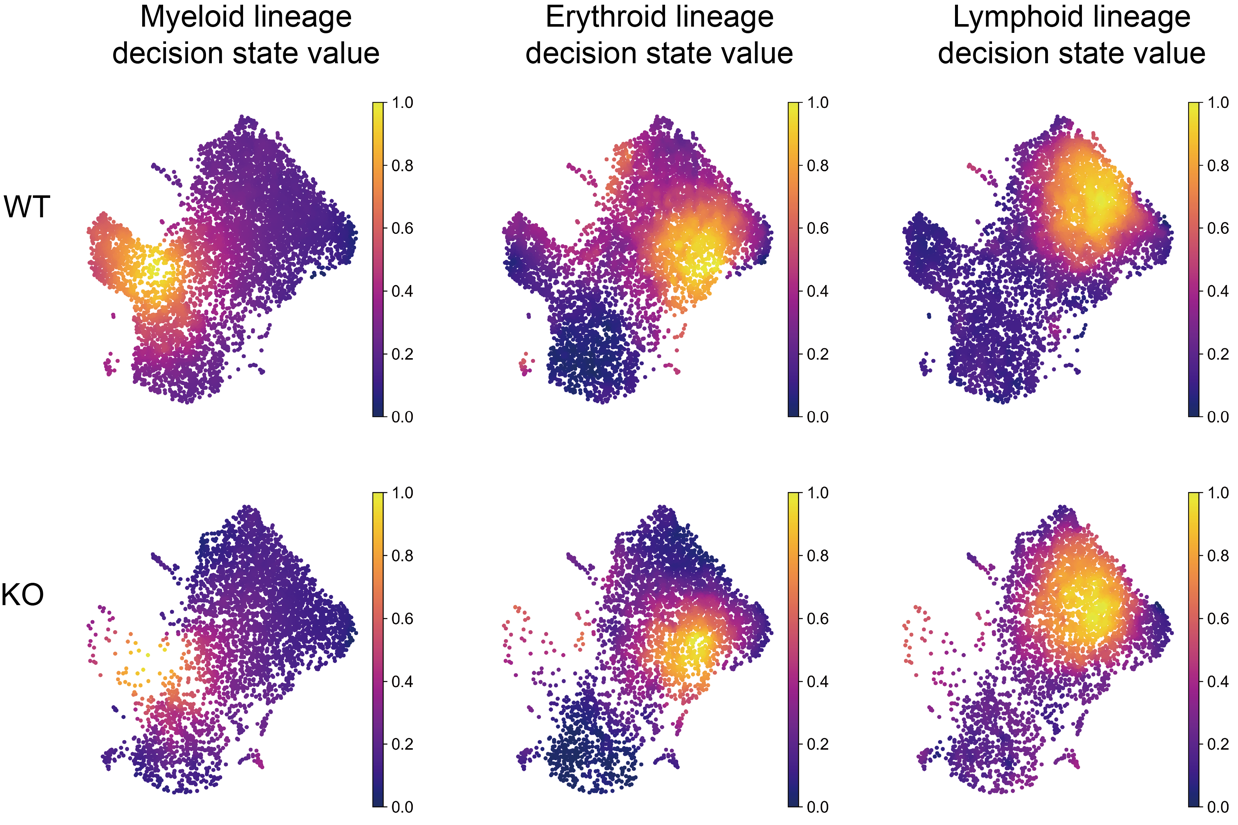

**Figure S11: Lineage decision value of Dapp1-Perturbed Datasets on UMAP**

The lineage decision intensity of myeloid, erythroid and lymphoid between Dapp1 knocked out and control groups.

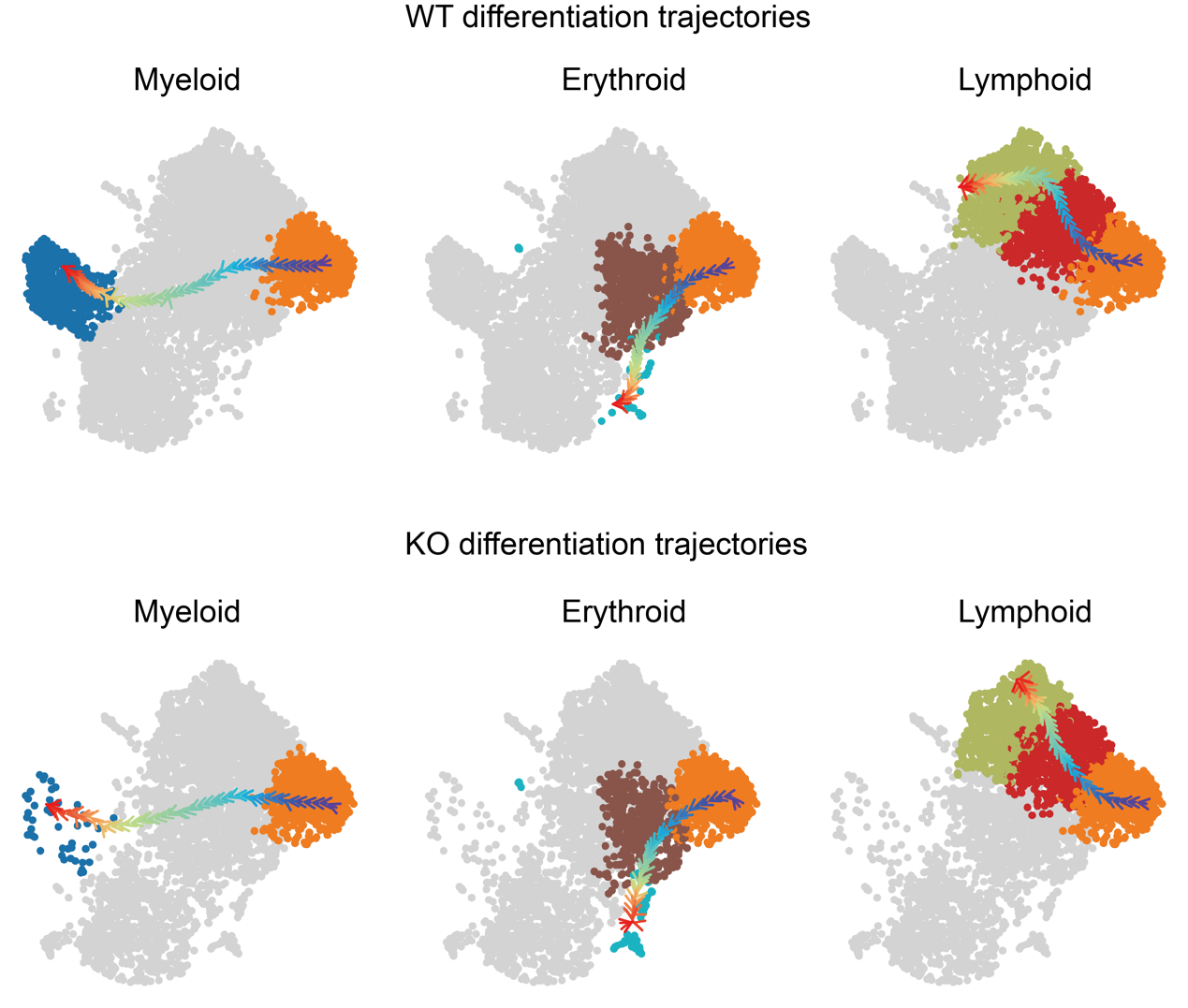

**Figure S12: Differentiation trajectories learned from lineage decision reward mode.**

The myeloid, erythroid and lymphoid differentiation trajectories of WT and KO group.

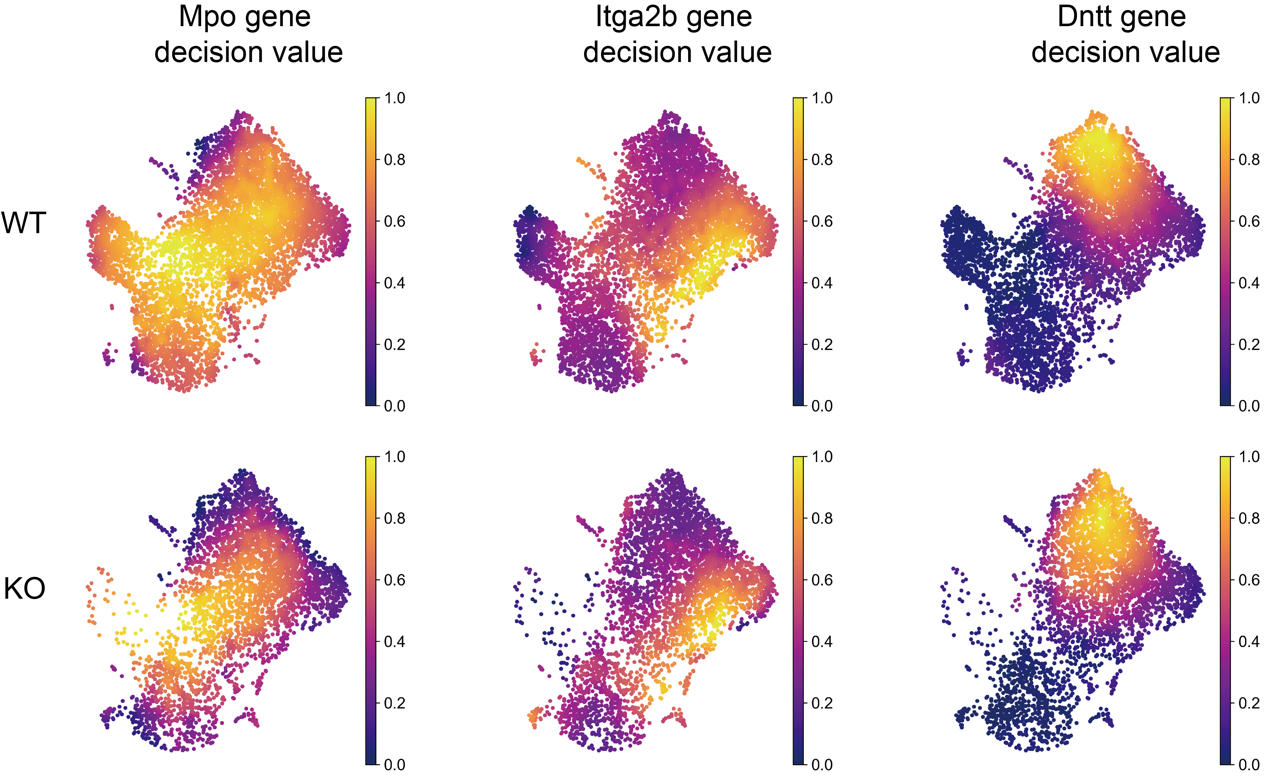

**Figure S13: Gene decision value of Dapp1-Perturbed Datasets on UMAP**

The gene decision intensity of Mpo, Itga2b and Dntt between Dapp1 knocked out and control groups.

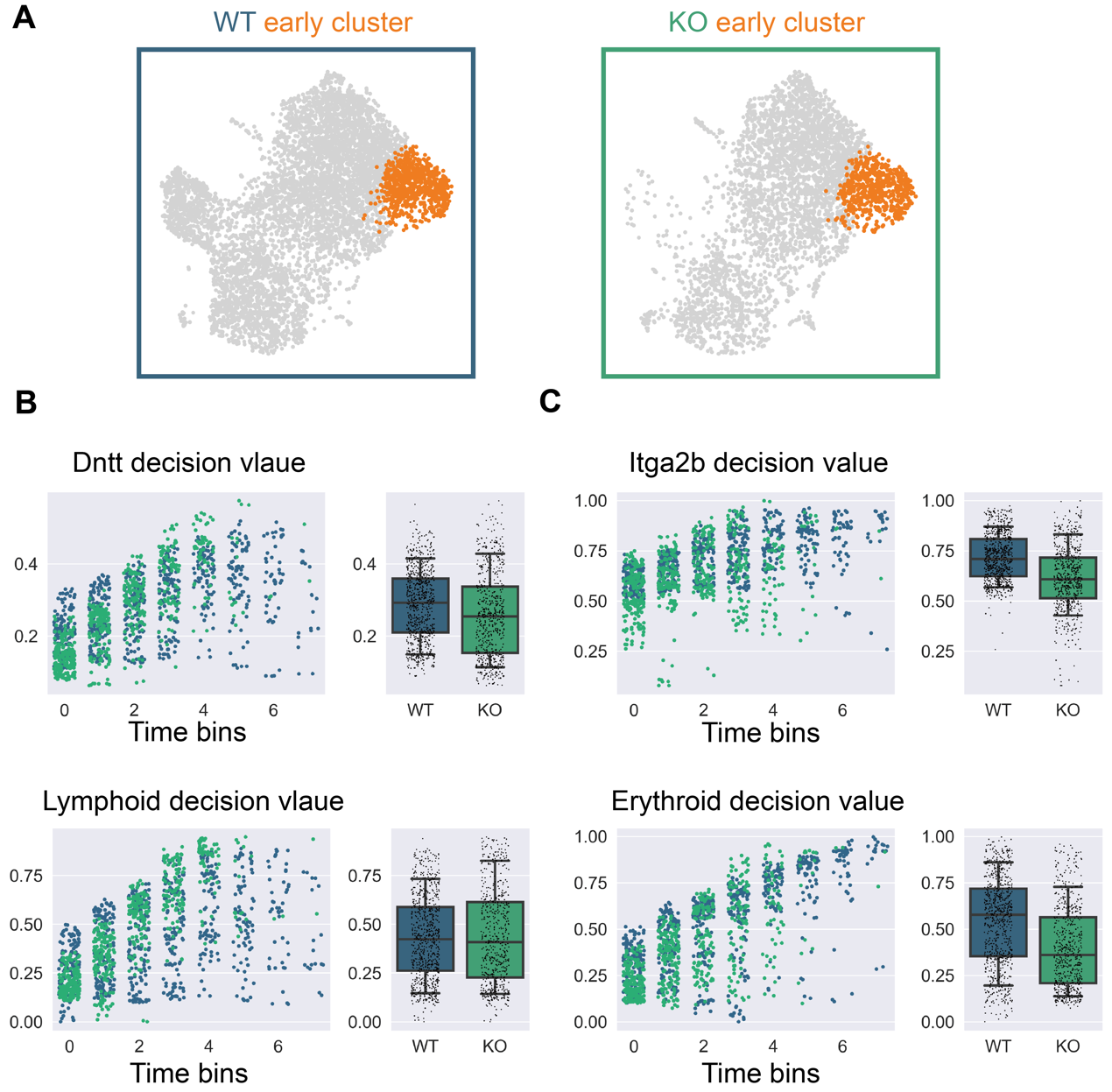

**Figure S14: Comparison of gene and lineage decision value within early cluster**

(A) The multipotent cluster of WT and KO group on UMAP. (B) The Dntt gene decision value and lymphoid lineage decision value along pseudotime within early cluster. (C) The Itga2b gene decision value and erythroid lineage decision value along pseudotime within early cluster.

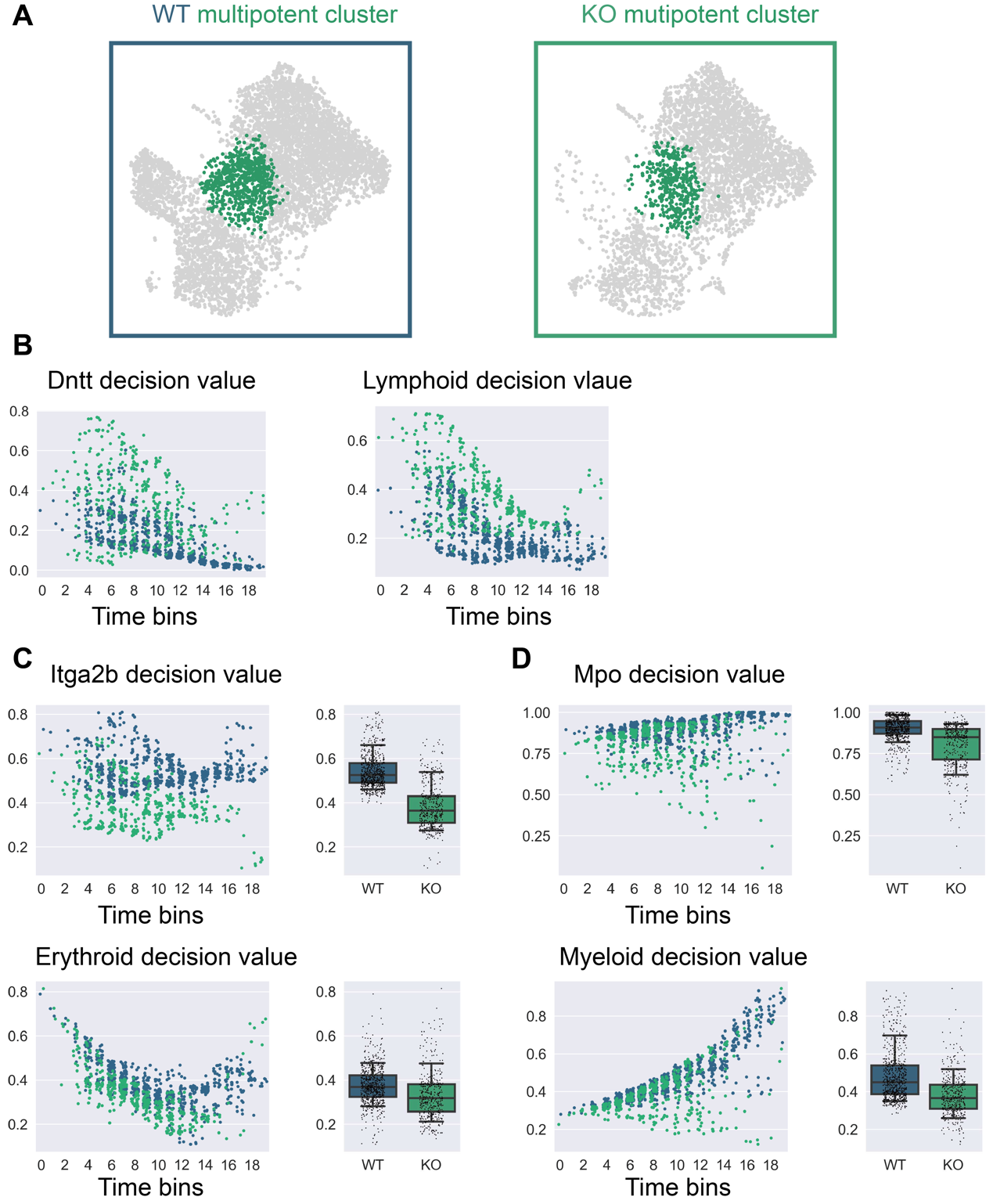

**Figure S15: Comparison of lineage and gene decision value within multipotent cluster**

(A) The multipotent cluster of WT and KO group on UMAP. (B) The Dntt gene decision value and lymphoid lineage decision value along pseudotime within multipotent cluster. (C) The Itga2b gene decision value and erythroid lineage decision value along pseudotime within multipotent cluster. (D) The Mpo gene decision value and myeloid lineage decision value along pseudotime within multipotent cluster.

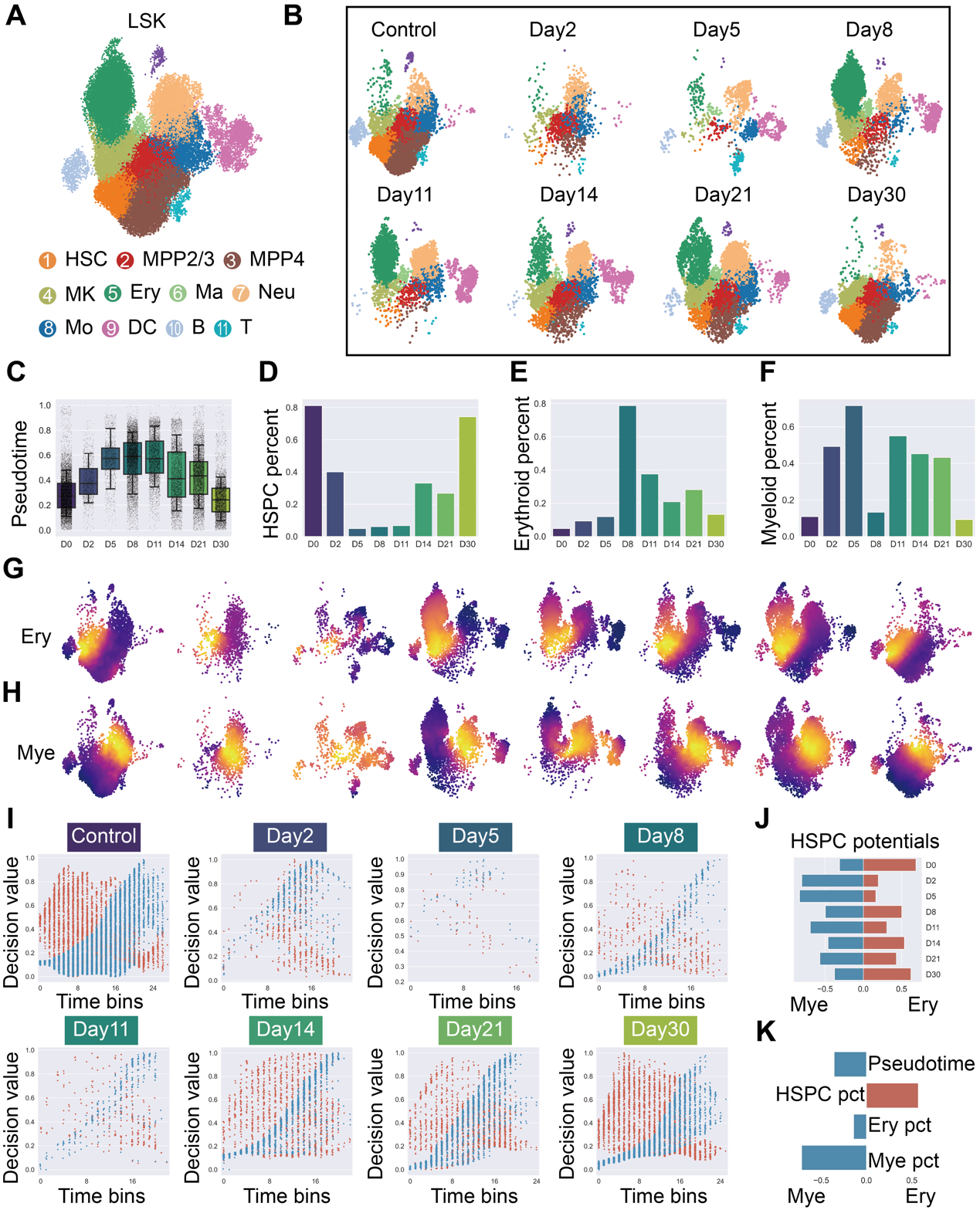

**Figure S16: Erythroid bias of primitive HSCs correlated with their proportion in time series of irradiation recovery**

(A) LSK (lin-Sca1+ckit+) phenotype bone marrow cells, clusters were annotated as HSC (Hematopoietic stem cell), MPP2/3(Multipotent progenitors 2/3), MPP4 (Multipotent progenitors 4), MK (Megakaryocyte), Ery (Erythrocyte), Ma (Mast cell), Neu (Neutrophil), Mo (Monocyte), DC (Dendritic cell), B (B cell), T (T cell) (B)Single cell atlas of irradiation injured LSKs at day points 2, 5, 8, 11, 14, 21, and 30, colored with annotated cell types in (A). (C) Box plot of pseudotime at each day points. (D) HSPC (HSC, MPP2/3, MPP4) proportion at each day points. (E) Erythroid cells (MK, Ery, Ma) proportion at each day points. (F)Myeloid cells (Gra, Mo, DC) proportion at each day points. (G) Erythroid lineage decision value projected on UMAP space at different time points. (H) Myeloid lineage decision value projected on UMAP space at different time points. (I) Lineage decision atlas for each time points within HSPC, erythroid decision value (dots colored in red) and myeloid decision value (dots colored in blue) along uniformly 50-binned pseudotime. (J) Horizonal bar plot of erythroid biased proportion (colored in red) and myeloid biased proportion (colored in blue) of HSPC at different time points. (K) Horizonal bar plot of the Pearson correlation of erythroid biased HSPC proportion, myeloid biased HSPC proportion at each time points with respect to the average pseudotime, HSPC proportion, erythroid proportion and myeloid proportion at each time points.

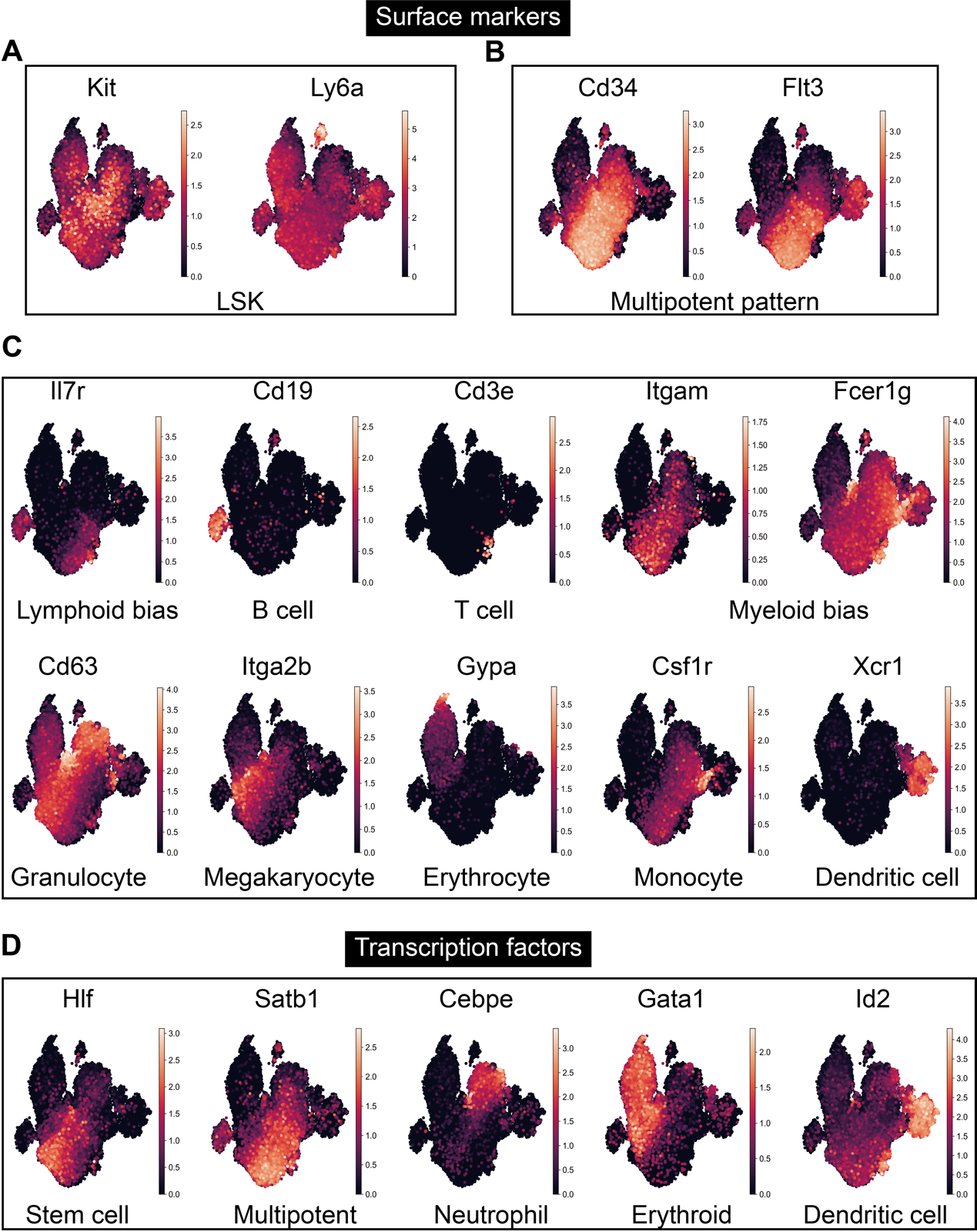

**Figure S17: Marker Genes for the Identification of Irradiated Hematopoietic Stem Cell (HSC) Types**

(A) Expression of Kit (c-Kit) and Ly6a (Sca-1) genes, serving as validation markers for the identification of Lineage-negative, Sca-1-positive, c-Kit-positive (LSK) cells. (B) Expression of Cd34 and Flt3, indicating the presence of multipotent progenitor components within the LSK population. (C) Key surface markers for the identification of various hematopoietic lineages, including Il7r for lymphoid bias, Cd19 for B cells, Cd3e for T cells, combined expression of Itgam and Fcer1g indicating myeloid bias, Cd63 for granulocyte bias, Itga2b for megakaryocytic bias, Gypa for erythrocytes, Csf1r for monocytes, and Xcr1 for dendritic cells. (D) Critical transcription factors that identify distinct hematopoietic lineages: Hlf for stem cells, Satb1 for multipotent progenitors, Cebpe for the neutrophil lineage, Gata1 for the erythroid lineage, and Id2 for the dendritic cell lineage.

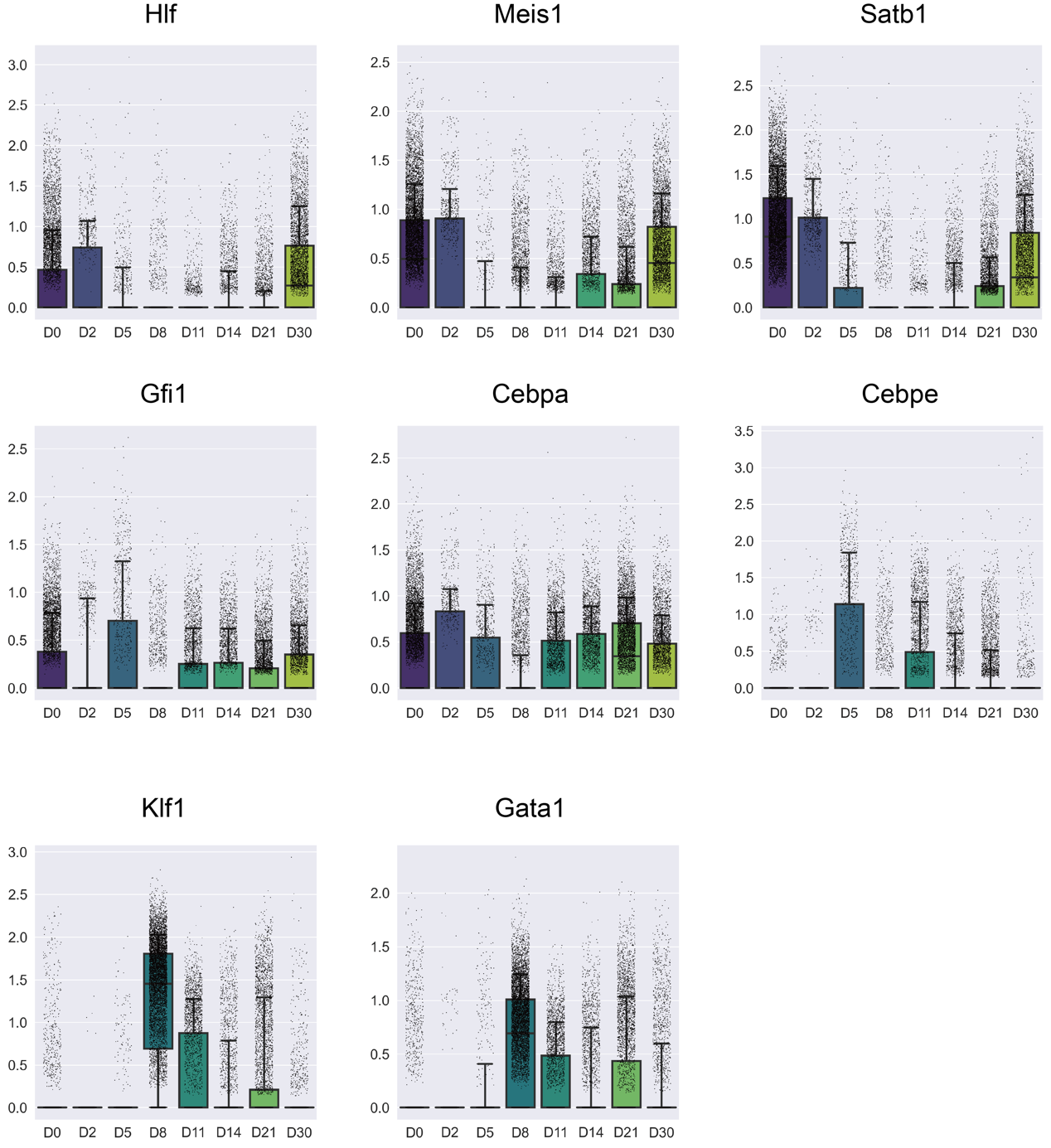

**Figure S18: Transcription factors patterns after irradiation**

(A) Hlf, Meis1, Satb1 and Hoxa9 were in similar expression pattern after irradiation, indicating the primitivity and multipotency of LSKs after irradiation. (B) Spi1, Gfi1, Cebpa and Cebpe were in similar expression pattern after irradiation indicating the myeloid bias potential of LSKs. (C) Klf1 and Gata1 were in similar expression patterns after irradiation indicating the erythroid bias potential of LSKs.

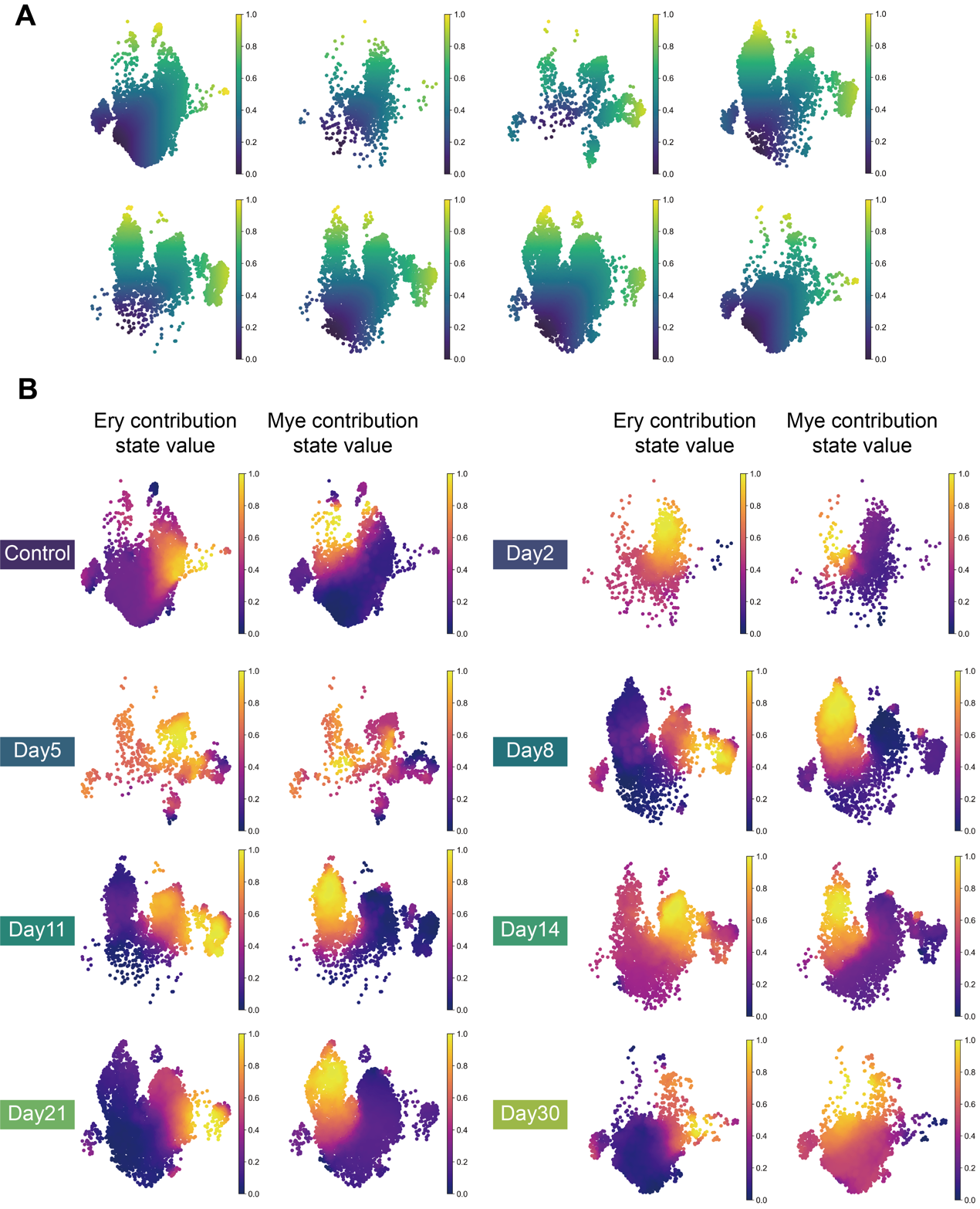

**Figure S19: Pseudotime and lineage contribution value of irradiation injured LSK**

(A) Pseudotime on different day points. (B)Erythroid and myeloid contribution value on different day points.

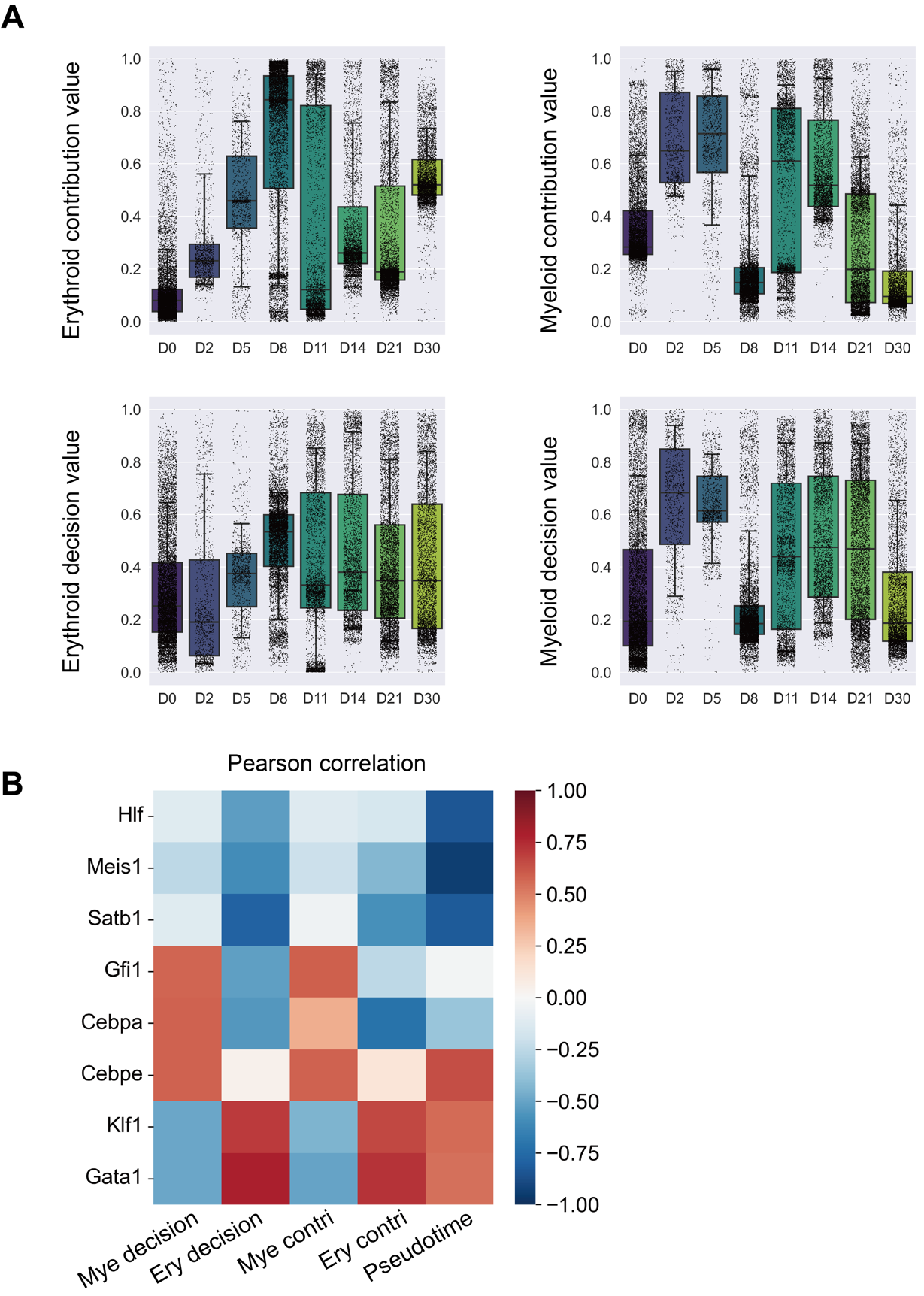

**Figure S20: Lineage Intensity of HSCs following Irradiation**

(A) This figure illustrates the intensity of erythroid and myeloid lineage contributions and decision value from normal LSKs and those at time points 2, 5, 8, 11, 14, 21, and 30 days post-irradiation. (B) Correlation heatmap of each time point’s lineage and decision value agianst gene expression values which including early genes Hlf, Meis1, Satb1, myeloid genes Gfi1, Cebpa, Cebpe and erythroid genes Klf1, Gata1.

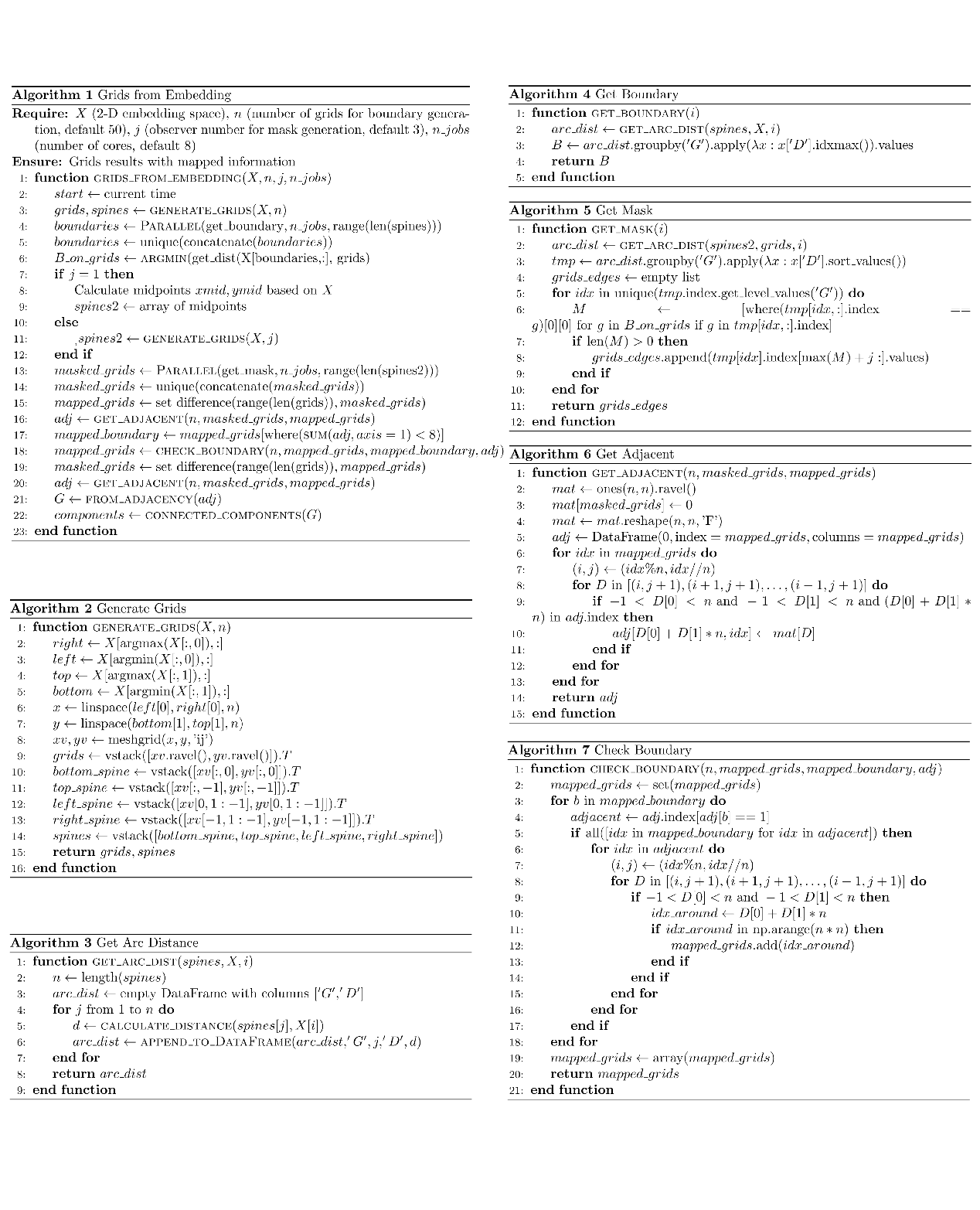

**Figure S21: Algorithms for generating grids embedding**

Assuming that most of the two-dimensional embeddings of large-scale single-cell data can represent the generation process and inherent dynamics of the data to some extent, a derivative grids representation of the embedding therefore can provide us with a more simplified and comprehensive perspective of the data space.

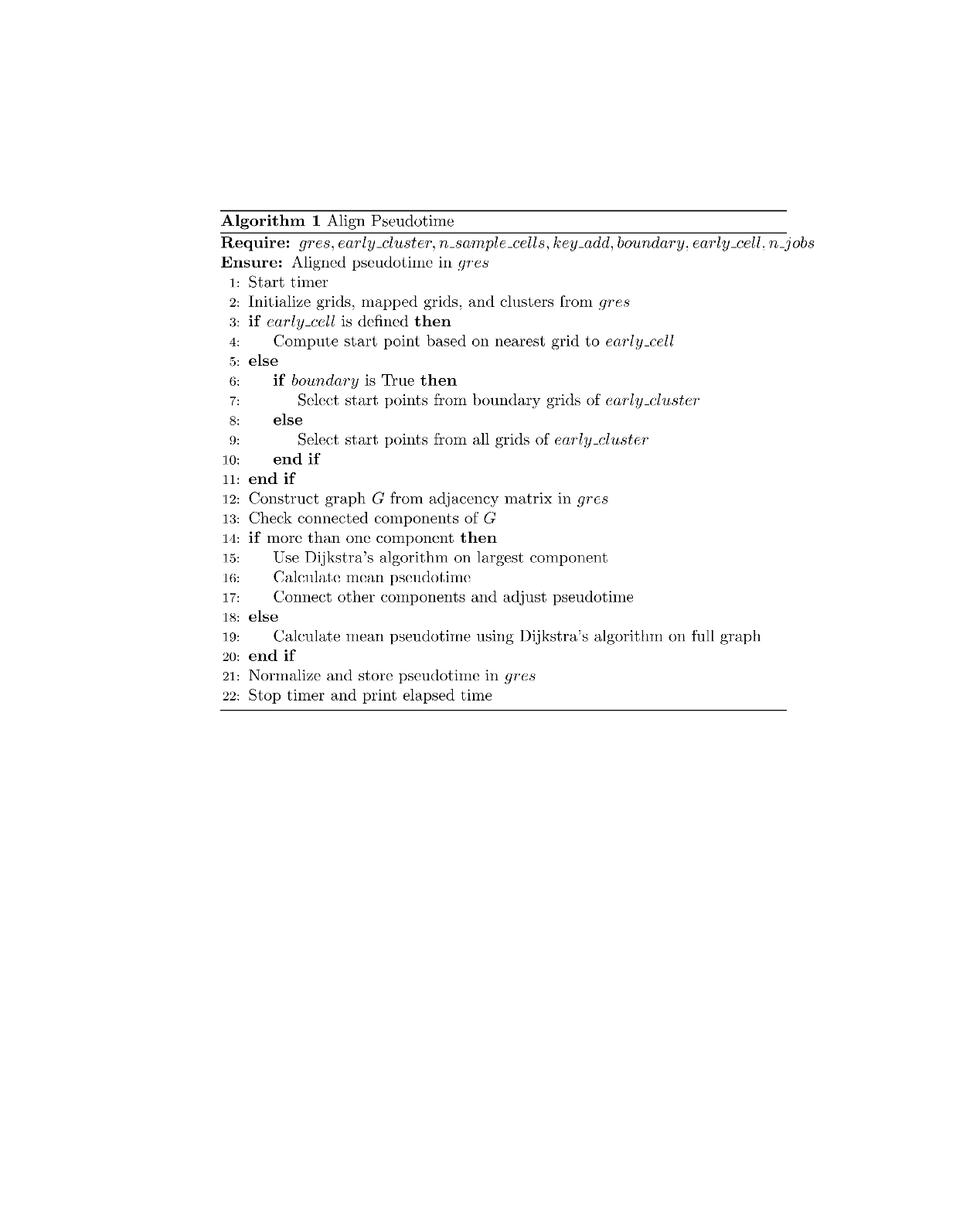

**Figure S22: Algorithm for pseudotime**

This function aligns pseudotime across a grid-based representation of data. It can start from a predefined early cell or a cluster, optionally restricting start points to grid boundaries. Pseudotime is calculated using Dijkstra's algorithm for shortest paths in a graph constructed from the grid adjacency matrix. The function handles multiple graph components by connecting them and adjusting pseudotime accordingly. The result is normalized and stored.

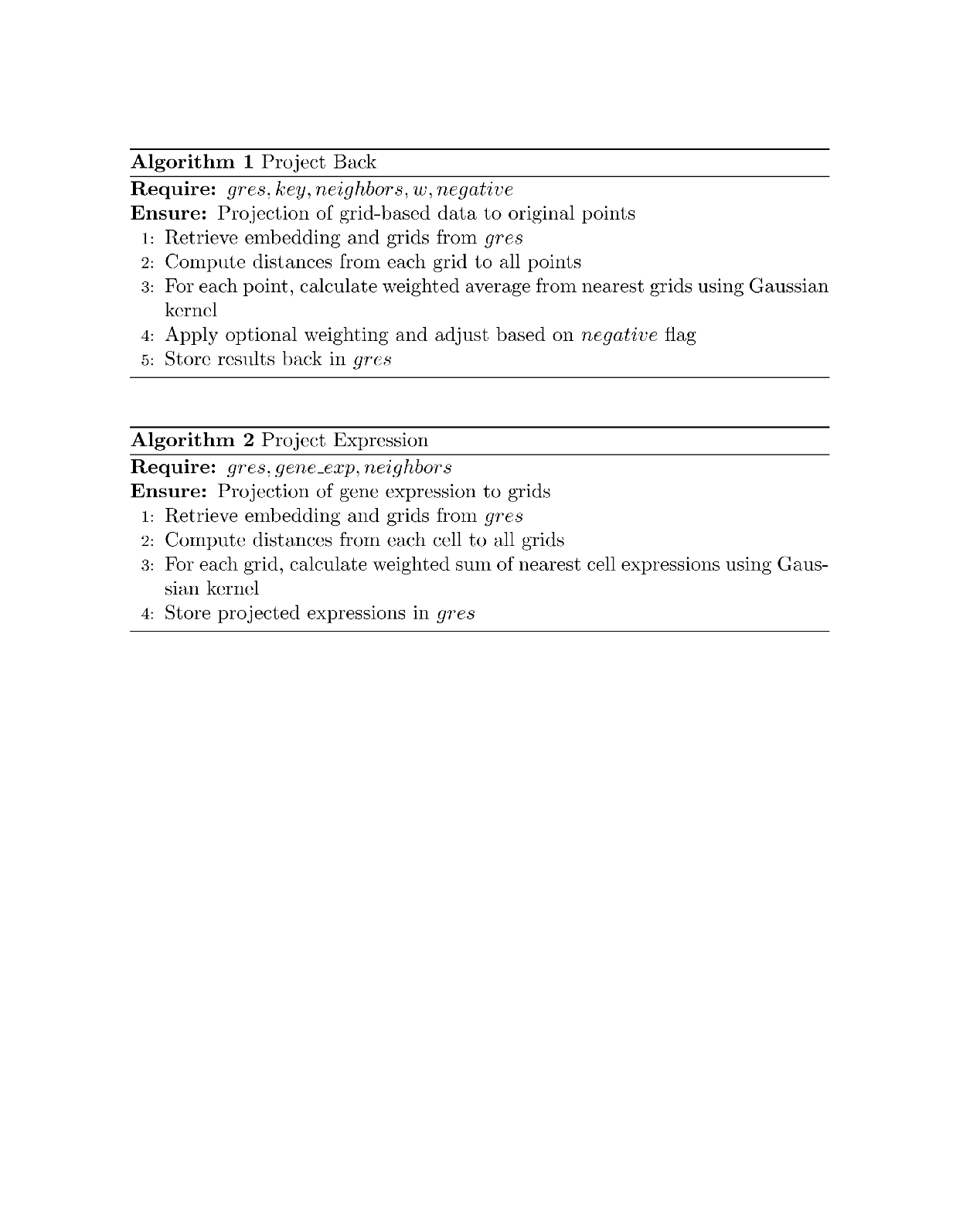

**Figure S23: Algorithms for projection**

Algorithm1 projects grid-based data back to the original points. It uses a Gaussian kernel to weigh the influence of the nearest grids on each point, optionally adjusted by a user-defined weighting factor. The results can be scaled to positive values and are stored back in the original data structure. Algorithm2 projects gene expression data from cells onto a grid-based representation. It calculates the weighted sum of the nearest cell expressions for each grid, using a Gaussian kernel to determine the weights. The projected expression data is then stored for further analysis.

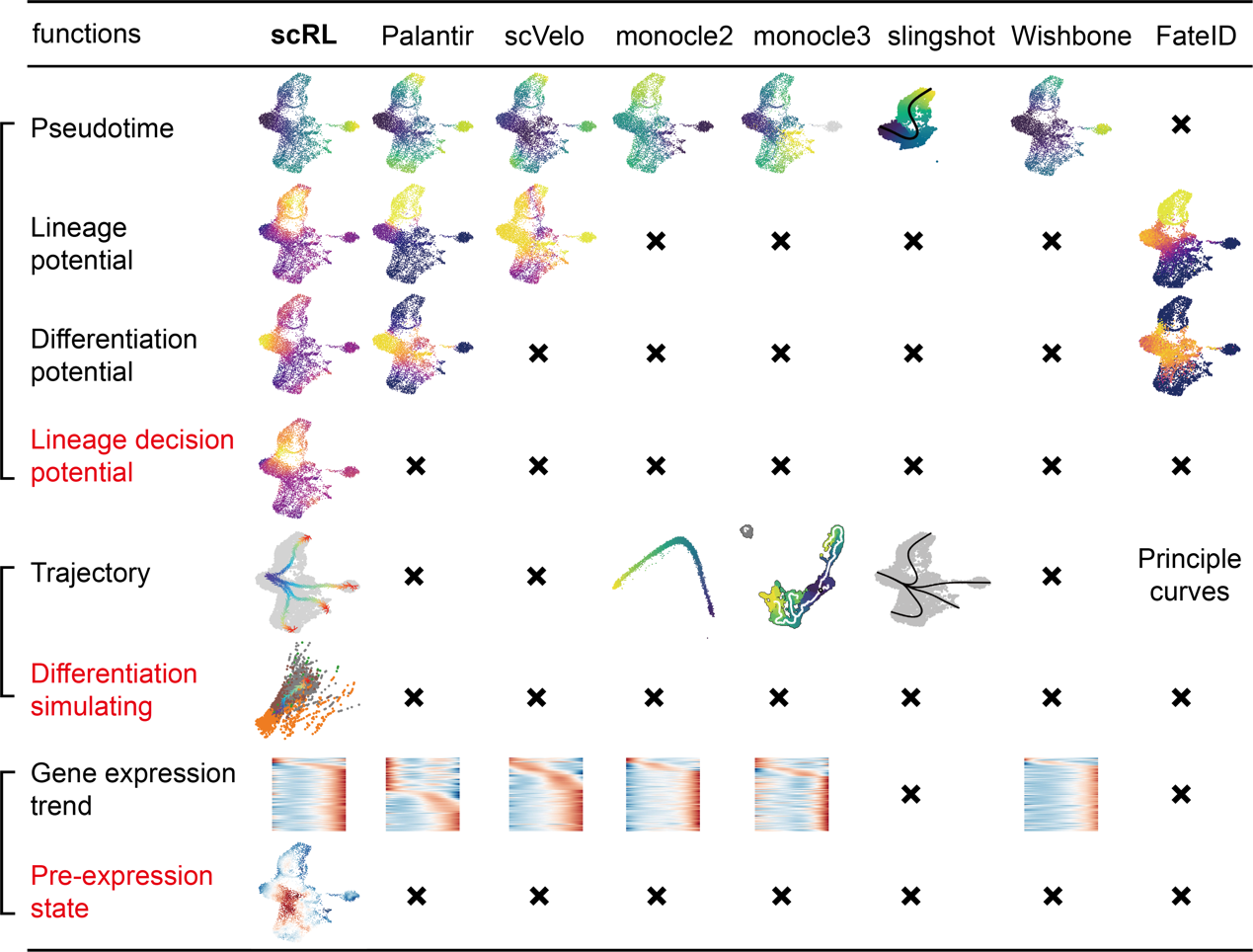

**Figure S24: Comparison between scRL and other methods**

A comparative analysis of scRL against state-of-the-art single cell pseudotime and differentiation analysis methods, including Palantir, scVelo, Monocle2, Monocle3, Slingshot, Wishbone and FateID. Functionalities of these methods were partitioned into three panels: pseudotime and potential, trajectory and simulating, and gene expression analysis. Red colored functions represent innovative contributions of scRL. Main results of each method are presented as scheme plots, and black crosses represent absent functions.

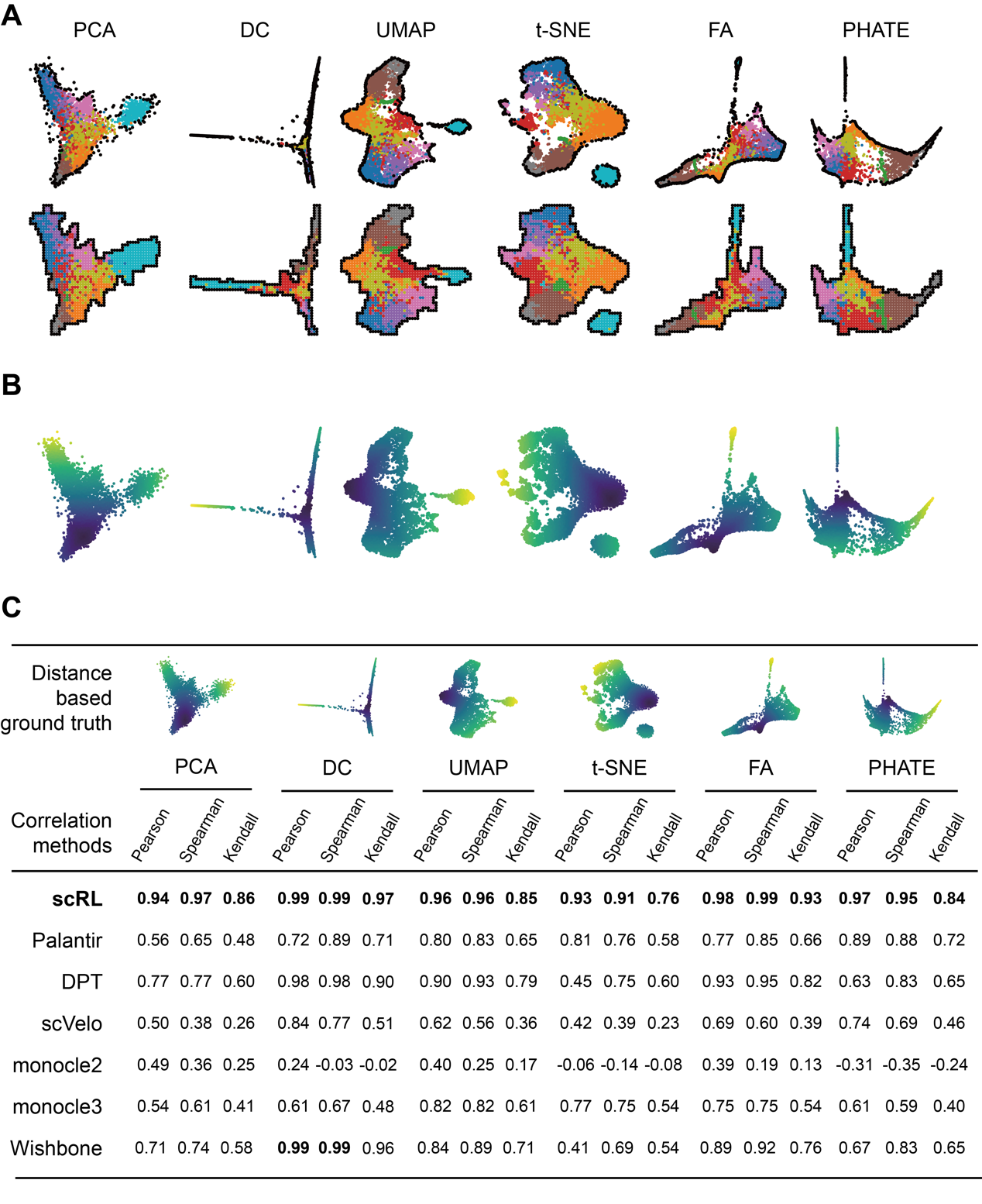

**Figure S25: Application of grid embedding and superiority of scRL’s pseudotime**

(A) Application of scRL’s grid embedding function to different dimensional reduction methods, black dots refer to the boundary points identified by scRL’s algorithm both in original embedding and the grid embedding. (B) Pseudotime inferred by scRL for different embeddings. (C) A comparison of correlation analysis between scRL’s pseudotime and other methods, scRL’s pseudotime was generated from different embeddings, and three kinds of correlation analysises were performed. The best score was in bold format.

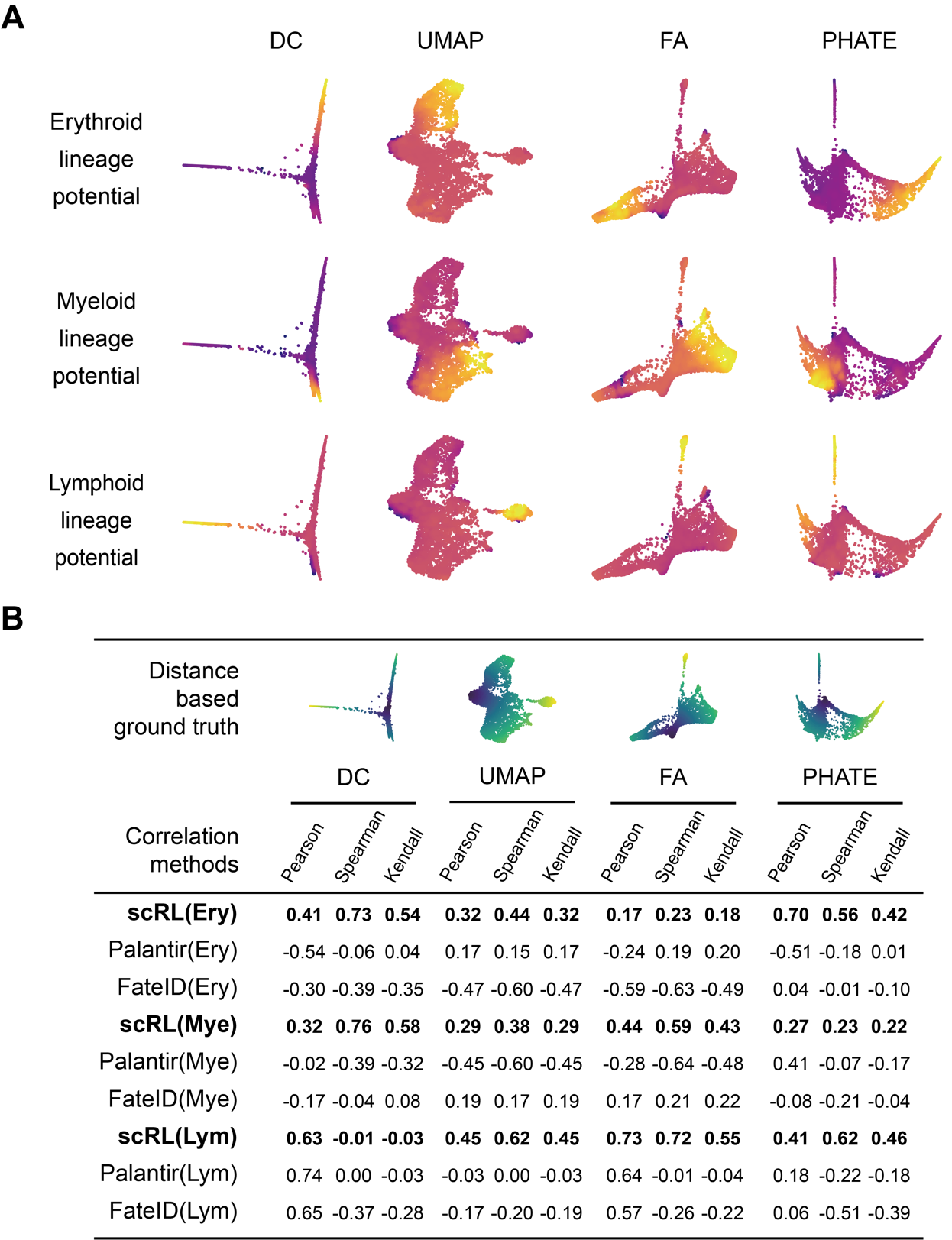

**Figure S26: Characters of scRL’s lineage potential compared with other methods**

(A) Erythroid, myeloid and lymphoid potentials recovered by scRL’s algorithm in different embeddings which DC is short for diffusion map components, UMAP is short for uniform mapping and projection, FA is short for force atlas and PHATE is name of the algorithm. (B) Comparison of correlation analysis between scRL’s lineage potential and Palantir as well as FateID. The score of scRL was in bold format.

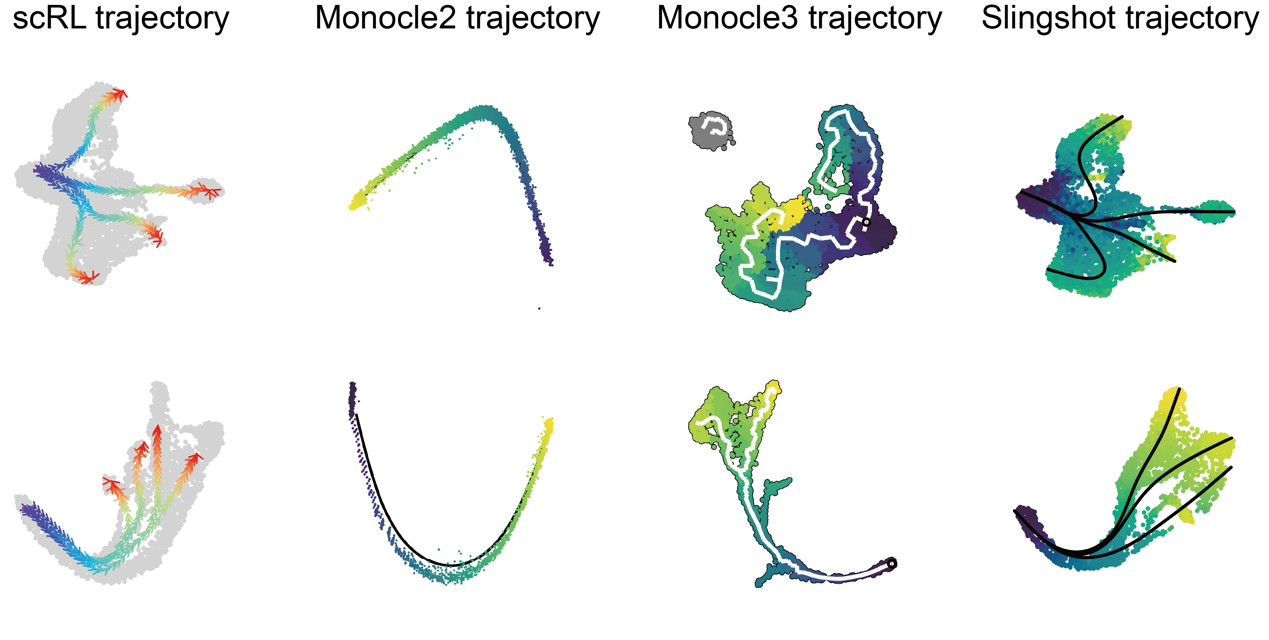

**Figure S27: Trajectories of scRL comparing with other trajectory methods**

A comparison of trajectories between scRL and monocle2, monocle3, slingshot trajectories based on the hematopoiesis and endocrinogenesis datasets.

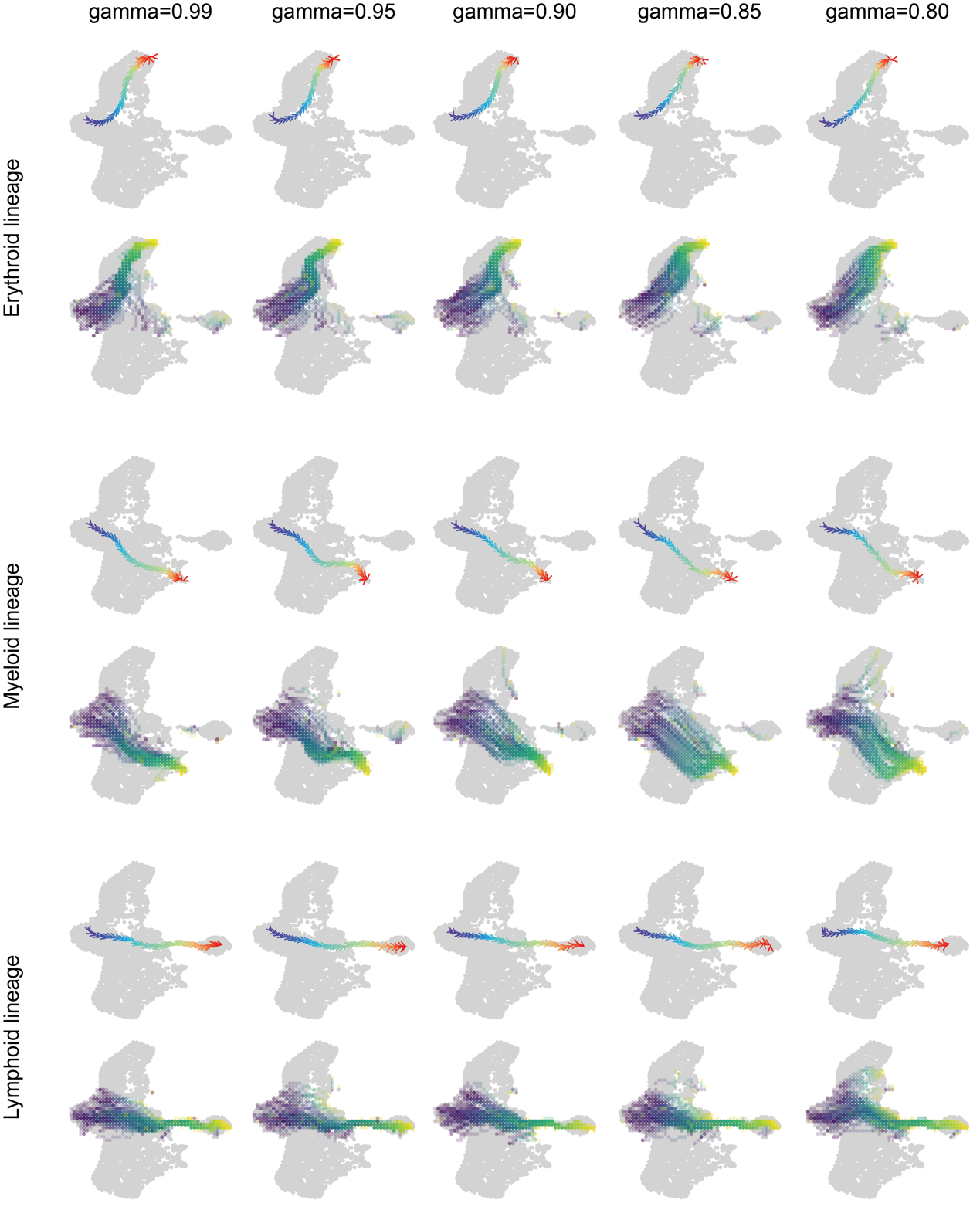

**Figure S28: Trajectories influenced by parameter gamma in Cd34+ bone marrow dataset**

Erythroid, myeloid and lymphoid lineages trajectories sampled based on gamma parameters of 0.99, 0.95, 0.9, 0.85 and 0.8.

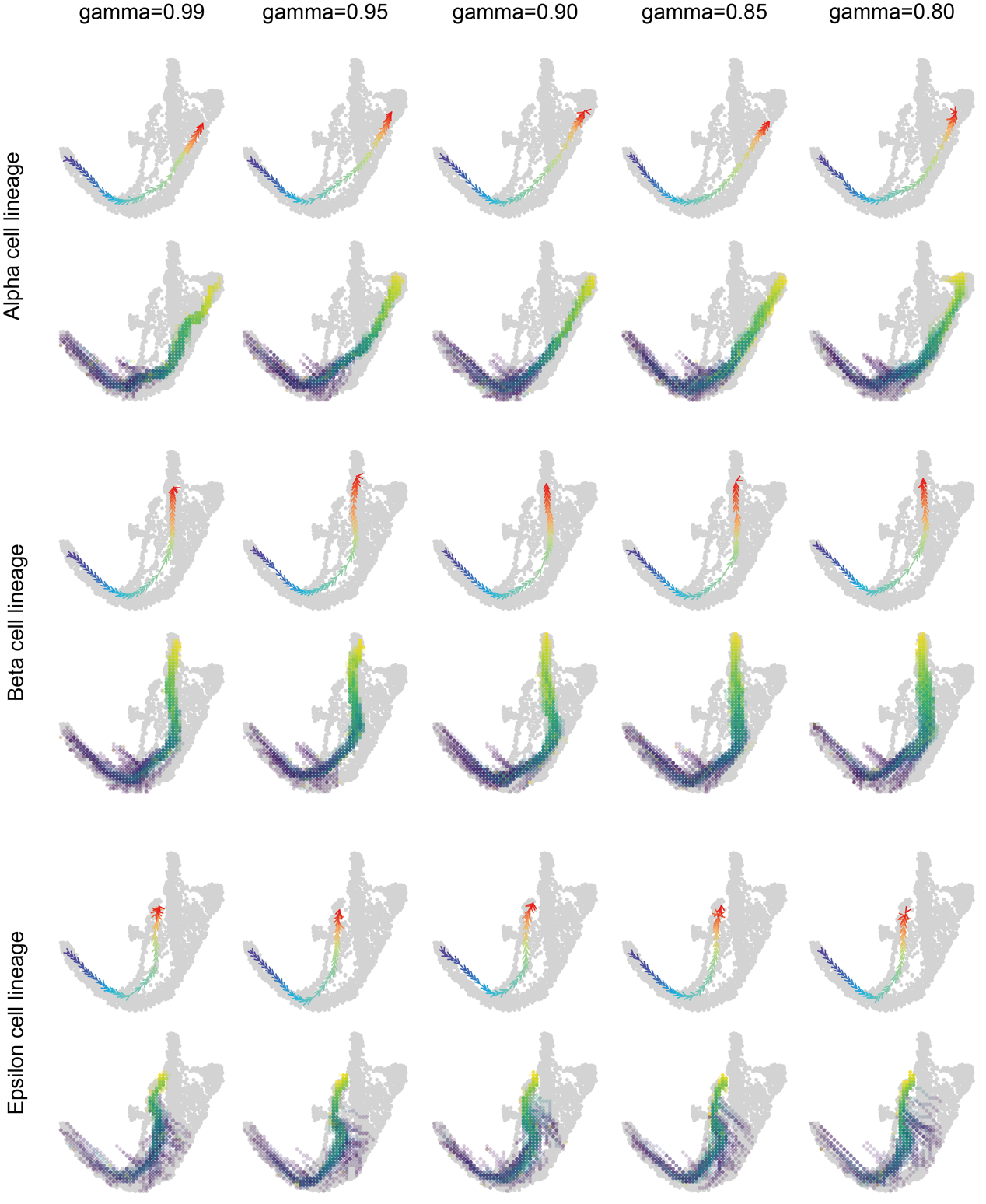

**Figure S29: Trajectories influenced by parameter gamma in endocrinogenesis dataset**

Alpha, beta and epsilon lineages trajectories sampled based on gamma parameters of 0.99, 0.95, 0.9, 0.85 and 0.8.

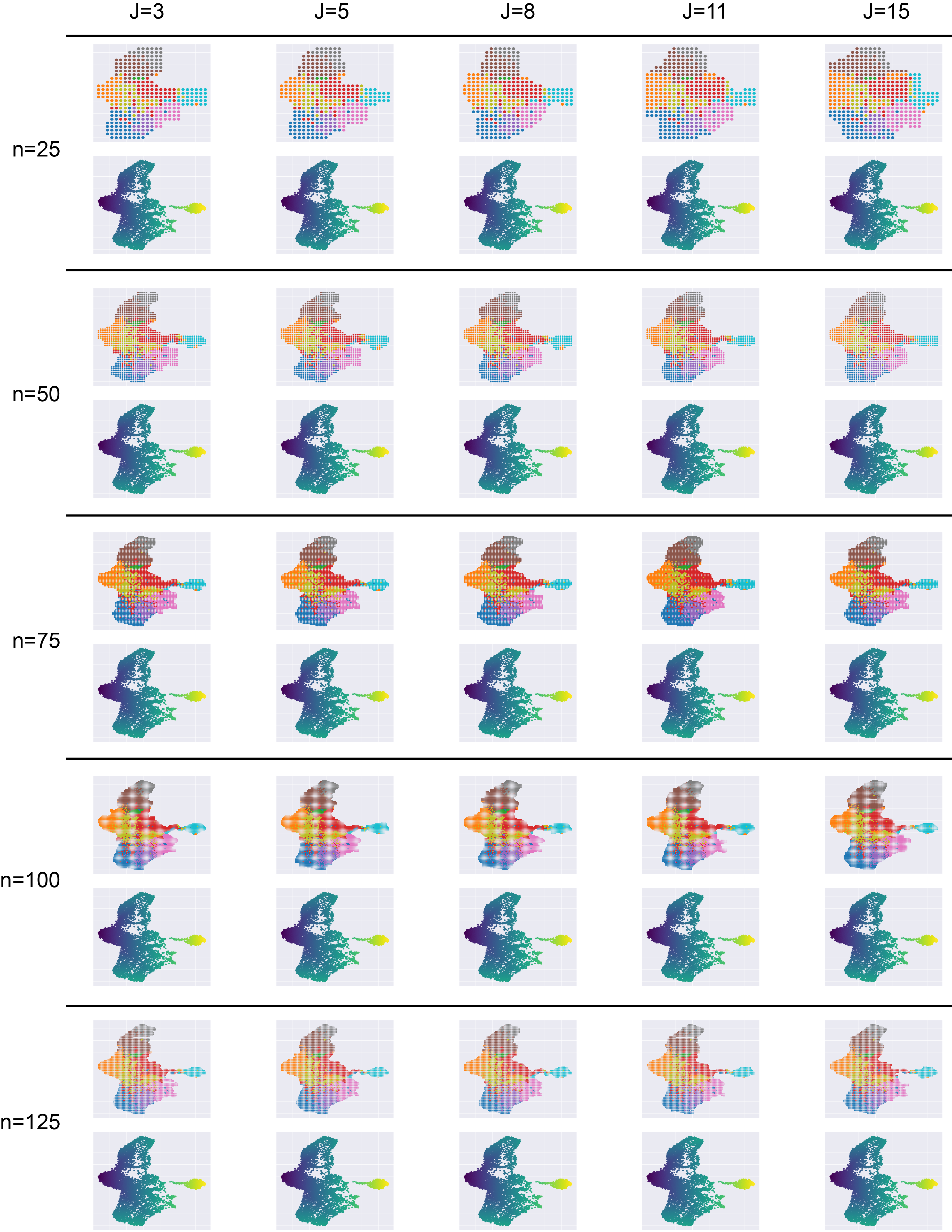

**Figure S30: Grid embedding function with different n and j on hematopoiesis dataset**

Application of different param n of grid embedding function	: 25, 50, 75, 100, 125 and for each param n with different param J: 3, 5, 8, 11, 15 on hematopoiesis dataset.

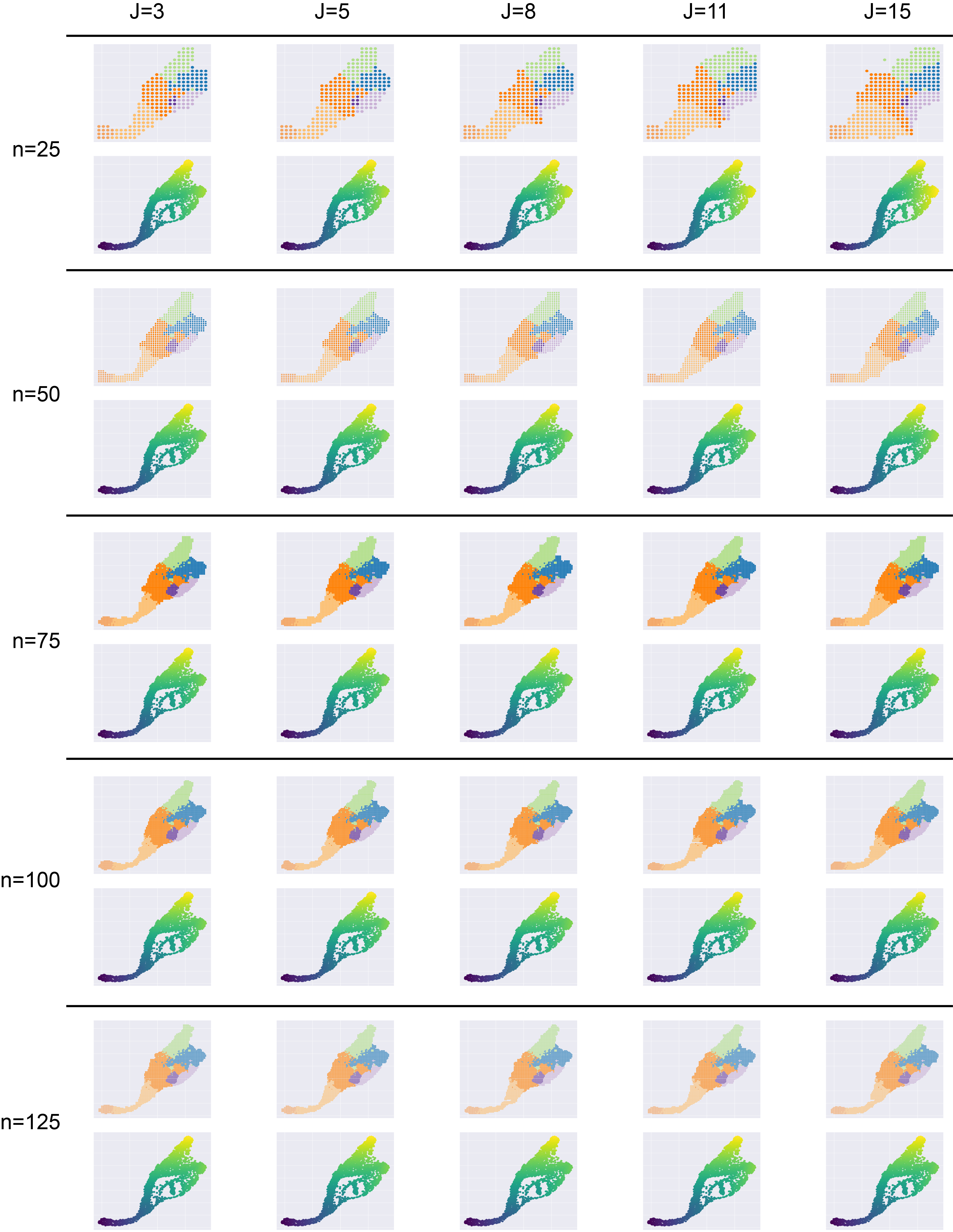

**Figure S31: Grid embedding function with different n and j on endocrinogenesis dataset**

Application of different param n of grid embedding function: 25, 50, 75, 100, 125 and for each param n with different param j: 3, 5, 8, 11, 15 on endocrinogenesis dataset

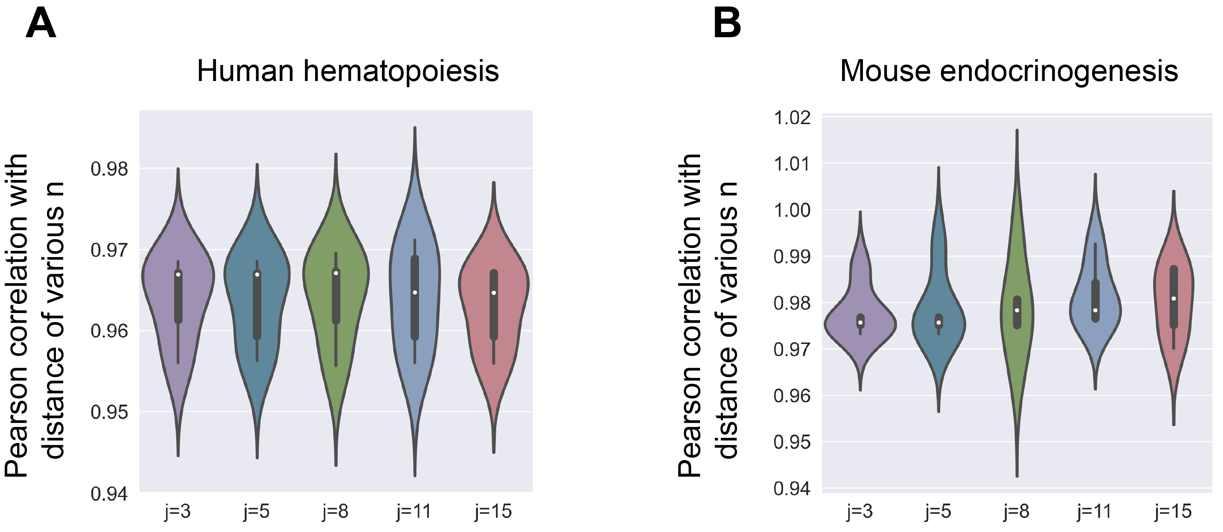

**Figure S32: Experiments Assessing the Pseudotime Alignment Function with Varying Parameters n and j**

(A) Violin plot depicting the pseudotime correlation scores obtained with varying values of parameter n for each fixed value of parameter j on the hematopoiesis dataset which allows us to assess the influence of both parameters on the accuracy and reproducibility of the pseudotime estimation. (B) Analogous to panel A, but on endocrinogenesis dataset, the pseudotime correlation scores obtained with varying values of parameter n for each fixed value of parameter j.
