## Supplementary materials for "scRL: Utilizing Reinforcement Learning to Evaluate Fate Decisions in Single-Cell Data"

**Pseudotime and lineage potential functions of scRL**

First, by employing a tabular Q-learning approach as an initial validation of the reinforcement learning environment, human hematopoietic CD34^+^ cell dataset with well-separated differentiation branches and mouse endocrinogenesis dataset with a stage-separated process underwent scRL’s grid embedding and pseudotime alignment functions within a UMAP 2-D space. The cluster information was subsequently projected onto the grid space (Fig. S1A). Meanwhile, specific rewards were generated for erythroid, myeloid, and lymphoid lineages within the grid embedding in the hematopoiesis dataset (Fig. S1B). For endocrinogenesis dataset, the rewards were set for early and late differentiation stages (Fig. S1B). Starting points were designated at HSC clusters for the hematopoiesis dataset and EP clusters for the endocrinogenesis dataset, with rewards setting at myeloid, erythroid, lymphoid, Fev^+^ cells, and mature cell clusters. Following the implementation of tabular Q-learning, we obtained and projected the final state values onto the original embedding (Fig. S1C). The results demonstrated that the training process was effective, evidenced by the convergence of both the cumulative reward obtained by the policy network and the maximum state value estimated by the critic network over time (Fig. S1D-E).

To evaluate the decoding capability of lineage decision states, we specifically used a dataset of human hematopoietic progenitor cells and a dataset of human CD34^+^ bone marrow cells (Fig. S2A, Fig. S3A). After aligning the pseudotime, we first conducted a comparative analysis between scRL and other prevalent methods such as Palantir, Wishbone, DPT, Monocle2 and Monocle3, based on the correlation coefficient with the Euclidean distance from the starting point. Notably, scRL was shown to achieve the highest score (Fig. S2B-C). Furthermore, by employing scRL's contribution reward mode, we assessed the commitment potentials of the primary MEP and GMP lineages properly (Fig. S2D-F). A similar trend was found in the erythroid and myeloid lineages of CD34^+^ bone marrow cells (Fig. S3B-F).

To evaluate the lineage decision state values, we performed an analysis on MEP and GMP lineages. The lineage decision values along pseudotime suggested that the decision values for MEP may peak during the intermediate stage of differentiation. Additionally, the contribution values reached their maximum at the most mature stage of differentiation. By comparing the weighted pseudotime for lineage decisions and the contribution strengths of MEP, we observed that the pseudotime weighted by decision value preceded the pseudotime weighted by contribution value (Fig. S2G-J). The trend is similar in GMP lineage and lineages of CD34^+^ bone marrow cells (Fig. S2K-N, Fig. S3G-N). These results indicated that the lineage decision values inferred by scRL represented a state prior to actual lineage commitment, highlighting the predictive capability of scRL in decoding lineage decisions. This finding underscored the utility of scRL in anticipating developmental outcomes in hematopoietic lineage commitment.

**Simulation function of scRL**

HSCs had the capability to differentiate into restricted progenitors including the erythroid and myeloid lineage. To simulate the differentiation of primary lineages in HSCs, we selected the HSC-MEP and HSC-GMP lineages from human phenotypic hematopoietic stem and progenitor cell (HSPC) dataset for differentiation simulation (Fig. S4A, Fig. S5A). During the exploratory training phase, an autoencoder was initialized at various starting points and progressed through 50 steps, inputting a single step and outputting the state difference. The simulation results demonstrated a correlation between the predicted pseudotime and branch information, revealing a gradual increase in the proportion of MEP and GMP lineages as the simulation advanced (Fig. S4B, Fig. S5B).

To validate the accuracy of the simulated paths, we compared the state space of the path points with the initial HSC targeting MEP and GMP lineages in the phenotypic HSC dataset. In the original space, the correlation with HSC decreased over pseudotime, while the correlation with MEP increased (Fig. S4C). Conversely, in the simulated space, the correlation coefficient between path points and HSC state spaces steadily declined as the paths progressed, while the correlation with MEP increased (Fig. S4D). Cluster heatmaps revealed that the correlation scores of HSC with MEP were higher in the simulated space than in the original space (Fig. S4E-F). Regarding gene expression levels, we observed an increase in correlation with MEP associated genes (Fig. S4G), while the correlation with HSC-associated genes decreased as pseudotime progressed. The trend was similarly observed in the GMP lineage (Fig. S5C-G). In the simulated gene expression space, the patterns mirrored those observed in the simulated lineage cells (Fig. S4H-J, Fig. S5H-J). We ranked the simulated genes based on their time correlation within the MEP lineage (Fig. S4K) and found that these genes exhibited lower time correlations in the original expression space for HSC-MEP (Fig. S4L). Moreover, the trends in simulated gene expression were more distinct for the HSC and MEP, compared to those observed in the original expression space (Fig. S4M-N). The results were consistent in the GMP lineage (Fig. S5K-N). In summary, these dynamic trends of correlation with HSC and various lineages illustrated the distinctive switching characteristics of fate decision in differentiation. These findings highlighted the complexity of gene expression dynamics and underscored the importance of simulation in understanding the nuanced changes across different hematopoietic lineages.

**Gene decision function of scRL**

scRL introduced a novel functionality in gene expression-based reward systems by enabling the identification of a pre-expression state for genes. Here, we utilized the datasets derived from human hematopoiesis and mouse endocrinogenesis to evaluate the effectiveness and convergence of scRL for gene expression analysis. We specifically selected marker genes indicative of distinct lineages: *GATA1* for the erythroid lineage, *IRF8* for the myeloid lineage, and *EBF1* for the lymphoid lineage. Additionally, we chose *Ngn3* as a marker for the early phases of endocrinogenesis and *Fev* to represent its intermediate stages^,^ (Fig. S6A). By employing these gene-specific markers, we generated rewards for grid embedding during the process of scRL (Fig. S6B).

Leveraging the tabular Q-learning algorithm, we explored the environment using an epsilon-greedy strategy to maximize gene rewards, while utilizing the Q-table to assess state values. The outputs were subsequently mapped back to the original embedding space (Fig. S6C). The meaningfulness of the training procedures was evidenced by the convergence of both episode rewards and peak state values over time (Fig. S6D-E). This convergence underscored the efficacy and robustness of our gene expression-driven reward system within the scRL framework, demonstrating its potential to enhance understanding of gene regulation dynamics in complex biological systems.

*GATA1* is a crucial transcription factor in MEP differentiation, while *CEBPA* is involved in the differentiation of the GMP lineage. To validate the pre-expression states predicted by scRL, we first observed that *GATA1* expression peaked at the mature stage in MEP lineage (Fig. S7A-B), and CEBPA expression in the GMP lineage exhibited a similar pattern (Fig. S7C-D). Following the scRL fate determination analysis, both *GATA1* and *CEBPA* displayed more primitive regions of high decision value intensity in the UMAP two-dimensional space compared to the contribution value (Fig. S7E-H).

As for *GATA1* in the erythroid lineage, the decision state value was shown to peak at the medium stage of differentiation, preceded the expression peak (Fig. S7I-J). The contribution state value for *GATA1* peaked at the mature stage of differentiation and demonstrated a stronger correlation with pseudotime in the MEP lineage than the original expression (Fig. S7K-L). Similarly, the decision state value and contribution state value of CEBPA also showed a similar trend in HSC-GMP lineage (Fig. S7M-P). These results suggested that the analysis of decision and contribution values provided a nuanced view of gene regulation during cell differentiation, offering insights into the timing and regulation of gene expression critical for effective lineage specification.

**scRL facilitates to discover the key regulators for lineage and cell fate decisions during HSCs differentiation**

To further illustrate the versatility of scRL, we tried to use it to discover the key regulators for lineage and cell fate decisions during hematopoietic stem cells (HSCs) differentiation. We initially mapped the natural differentiation trajectory of LSK (Lin^-^Sca1^+^ckit^+^) cells using PALANTIR diffusion and t-SNE embedding spaces. Then, we applied scRL to create a grid embedding, anchored by the primitive genes *Hlf* and *Mllt3*, and analyzed the dynamic process of cell differentiation through pseudotime and velocity vector fields. This approach enabled us to identify rapidly upregulated genes during early lineage differentiation, including *Dapp1*, *Adgrl4*, *Lmo2*, *Cbfa2t3*, *Malat1*, *Ctla2a* and *Hlf*, with *Dapp1* showing a strong temporal association (Fig. S8A). Further analysis using DDRTree traced the differentiation trajectory to predict the lineage fate decision statuses (Fig. S9A). By intersecting dynamically expressed genes with those identified as significant by BEAM, we focused on 280 genes enriched in hematopoietic differentiation-related Gene Ontology Biological Process (GOBP) terms (Fig. S9B-C). Additionally, we conducted a gene association analysis using LSK datasets with representative lineage markers, *Dach1* for myeloid lineage differentiation and *Dntt* for lymphoid lineage differentiation. Intriguingly, *Dapp1* emerged as highly correlated with *Dach1* and exhibited a lesser correlation with *Dntt* (Fig. S9D-E). The intersecting genes from these datasets further substantiate the close association of *Dapp1* with myeloid differentiation within the LSK population (Fig. S9F).

To validate whether scRL can effectively capture the impact of perturbation in key dynamical genes on hematopoietic stem cell differentiation states, a single-cell atlas was constructed from the bone marrow LSK cells with conditional *Dapp1* knockout in the hematopoietic compartment (Fig. S8B). The lineages within this atlas were annotated based on the expression patterns of key transcription factors (Fig. S10). This mapping revealed a significant disruption in lineage commitment homeostasis, characterized by a marked reduction in the myeloid lineage (Fig. S8C). Additionally, pseudotime and expression of marker genes *Cd34*, *Mpo* and *Ctsg* suggesting an impediment to the cellular maturation process (Fig. S8D).

Utilizing the myeloid, lymphoid, and erythroid clusters as a basis for reinforcement learning, we observed shifts in lineage decision values (Fig. S11). Analysis of lineage decision trajectories along pseudotime revealed that *Dapp1* deficiency led to significant perturbations (Fig. S8F). Specifically, the decision values for myeloid and erythroid lineages decreased, while those for the lymphoid lineage increase, indicating an imbalance in fate decision after *Dapp1* was conditional knockout. However, the differentiation trajectories learned were not sufficiently distinct (Fig. S12). To further validate these findings, we analyzed gene decision values specific to each lineage: *Mpo* for myeloid, *Itga2b* for erythroid, and *Dntt* for lymphoid (Fig. S13). The trajectories of these gene decision values underwent notable changes (Fig. S8G). Moreover, *Dapp1* deletion in hematopoiesis resulted in a decrease in the decision values for the myeloid gene *Mpo* and the erythroid gene *Itga2b*, while the decision value for the lymphoid gene *Dntt* increased (Fig. S8H).

Focusing on the earliest cluster (Fig. S14A), we observed that both the decision trajectory for the myeloid gene *Mpo* and the myeloid lineage decision trajectory are disrupted if *Dapp1* was conditional knockout (Fig. S8I). Specifically, both gene and lineage decision values for myeloid decrease after the knockout (Fig. S8K). In contrast, the lymphoid lineage decision trajectory and the *Dntt* gene decision value did not decrease (Fig. S14B). While the erythroid lineage and *Itga2b* gene decision values also decreased, this change was less pronounced compared to that observed in the myeloid lineage and *Mpo* (Fig. S14C). In the earliest cluster, lymphoid potential remained largely unchanged. However, within the multipotent cluster (Fig. S15A), both lymphoid and *Dntt* decision values are enhanced (Fig. S8L), along with their decision trajectories along pseudotime (Fig. S15B). Additionally, we note a decrease in *Itga2b* and erythroid decision potential (Fig. S15C), as well as in *Mpo* and myeloid decision values (Fig. S15D). The application of scRL successfully addressed the challenge of assessing lineage fate decisions in single-cell data. Moreover, the integration of scRL with single-cell subpopulation differentiation provided a comparative analysis of lineage fate decisions across different differentiation levels, our results were confirmed with single-cell data from key differentiation gene perturbations.

**scRL decodes the fate decision atlas of HSCs after irradiation**

After validating the effectiveness of scRL, we sought to apply it to analyze the fate decision atlas of HSCs under pathological conditions. As is known, hematopoietic cells are sensitive to ionizing radiation, while the cell fate transitions of HSCs after radiation injury remain largely unknown. Here, we first collected LSK cells from mice inflicted with total body ionizing radiation at different days after radiation and performed single-cell sequencing (Fig. S16A). Then, a comprehensive fate decision atlas of HSCs post radiation exposure was constructed with scRL. Eleven biologically significant clusters were marked by critical gene expressions, consistent with previous studies (Fig. S17A-D). The number of HSPC, especially HSCs, sharply reduced after radiation and reached the bottom around day 5, followed by stepwise recovery. Meanwhile, an erythroid commitment bias emerged after radiation, peaking at day 8, and persisted until day 21(Fig. S16B).

Utilizing pseudotime as a metric for cellular primitivity, the recovery process was interpreted as HSCs regaining their primitive state (Fig. S16C). As shown, HSPC components decreased during the injury phase but recovered by day 30, correlating with the expression patterns of *Hlf*, *Meis1*, and *Satb1* (Fig. S16D, Fig. S18A). However, both erythroid and myeloid components increased before day 8 after radiation (Fig. S16E), which corresponded to the expression patterns of *Klf1* and *Gata1*, as well as the expression patterns of *Gfi1*, *Cebpa*, and *Cebpe* respectively (Fig. S16F, Fig. S18B-C).

To decipher the fate decisions of erythroid and myeloid lineages following irradiation, we devised lineage-specific reward environments and computed pseudotime for each day post-irradiation (Fig. S19A). By employing lineage rewards, we utilized the critic network to assess lineage decision values for both erythroid and myeloid lineages (Fig. S16G-H). Additionally, we determined the lineage contribution values for both erythroid and myeloid lineages (Fig. S19B, Fig. S20A-B). We then conducted a comparative analysis of erythroid (colored in red) and myeloid (colored in blue) lineage decision values across pseudotime within selected HSPC subpopulations (Fig. S16I). Although the fate decision potential of erythroid-biased stem cells decreased dramatically after radiation, it rapidly returned to near normal level at day 8. Conversely, the proportion of myeloid-biased stem cells increased following irradiation injury and subsequently decreased as the erythroid-biased stem cells increased (Fig. S16I-J). Interestingly, the erythroid potential of HSPCs showed the strongest correlation with the overall proportion of HSPCs, whereas the myeloid potential was closely associated with the proportion of myeloid cells during the early days after radiation (Fig. S16K). These findings indicate that the surviving HSCs after radiation preferentially differentiate into erythroid lineage cells, and the increase of myeloid potential is possibly ascribed to the radiation resistance property of myeloid progenitor cells. In fact, the priority of erythroid fate decision occurs not only post irradiation exposure but also under physiological status, probably due to the indispensable role of erythrocytes in the body. Therefore, integrating reinforcement learning with single-cell analysis can reveal deeper insights into HSCs fate decisions under physio pathological conditions.

**The Significance of Transition Cells and Reinforcement Learning in Deciphering Cell Fate Decisions**

Transition cells play a crucial role in understanding cell fate decisions, as they reflect the transient dynamics during a cell-fate switch and exhibit mixed identities from multiple cell states. These characteristics make transition cells particularly interesting for unraveling the complex mechanisms of cell fate decisions and understanding how cells respond to various pathophysiological stimuli. Identifying transition states and critical transition signals during cell differentiation in single-cell data is essential for gaining deeper insights into these processes.

The application of reinforcement learning in single-cell data analysis can be viewed through the dual processes of exploration and exploitation, akin to navigating an intricate map of cellular differentiation. The algorithm begins with random exploration to uncover unknown patterns and pathways within the complex differentiation landscape. As it progresses, the exploration is refined into a strategic approach, evaluating each decision point or node for its potential direction in cellular differentiation. The algorithm identifies and moves towards reward nodes, representing optimal differentiation pathways or significant biological states. Consistently successful strategies become reinforced, shaping a more deterministic and efficient path through the differentiation map. The goal is to efficiently guide the agent from an initial state to desired lineage outcomes, optimizing the trajectory based on accumulated knowledge and rewards.

UMAP serves as a critical bridge, connecting sophisticated computational analysis with accessible and interpretable visual representations, facilitating a deeper understanding of the dynamic processes governing cell fate decisions. UMAP's flexibility in preserving the geometric structure of the data and its scalability make it a superior choice for visualizing cell differentiation, where lineage subpopulations emerge as targeted directions in the maturation process.

In conclusion, studying transition cells and applying reinforcement learning techniques, in combination with visualization methods like UMAP, can provide valuable insights into the complex mechanisms of cell fate decisions and help decipher the intricate landscape of cellular differentiation.

**Analysis of human CD34^+^ bone marrow dataset**

The dataset, containing 5780 cells, was downloaded from scVelo. Preprocessing was conducted using Scanpy, where cells were normalized based on total gene counts. From this, 2000 highly variable genes were selected for subsequent principal component analysis. Default parameters were used to calculate the neighbor graph and UMAP. Cell types were adopted from the original study. Utilizing scRL with default settings$\left( n=50,j=3 \right)$, a grid embedding was created. The erythroid lineage encompassed Ery_1, Ery_2, and Mega clusters, while the myeloid lineage encompassed Mono_1, Mono_2, and DCs. The lymphoid lineage encompassed the CLP cluster, which were designated as rewards, with HSC_1 set as the starting cluster. Gamma parameters of 0.9 and 0.8 were used for decision reward and contribution reward modes, respectively. The model underwent training across 5000 episodes.

**Analysis of mouse endocrinogenesis dataset**

The dataset, encompassing 2531 cells, was sourced from scVelo. Preprocessing via Scanpy involved normalizing cells by total gene counts. Out of this, 1000 highly variable genes were chosen for downstream principal component analysis. Default settings were applied for calculating the neighbor graph and UMAP, with cell annotations taken from the original study. scRL was utilized to generate a grid embedding with parameters$\left( n=50,j=3 \right)$. The reward lineages encompassed alpha, beta, delta, and epsilon clusters, while Ngn3 low, Ngn3 high, and Fev+ were set as starting clusters. Gamma values of 0.9 and 0.8 were chosen for decision reward and contribution reward modes, respectively. The model was trained over 5000 episodes.

**Analysis of human phenotypical HSC dataset**

The dataset was publicly accessible via GSE117498. It encompassed HSC, MPP, MLP, CMP, MEP, GMP and PreBNK, from which 5140 highly variable genes were chosen for downstream PCA, neighbor graph construction, and UMAP using default settings. Batch information served as cell types. A grid embedding was created using scRL with default parameters$\left( n=50,j=3 \right)$. MEP and GMP were designated as reward clusters, with HSC as the starting cluster. Gamma values of 0.9 and 0.8 were set for decision reward and contribution reward modes, respectively。

**Analysis of human AML dataset**

The dataset, containing 94311 cells, was publicly available through GSE185993. Cells were normalized by total gene counts, and 2000 highly variable genes were chosen for downstream analysis. The neighbor graph was generated using the top 30 principal components, considering 30 neighbors. UMAP and Leiden clustering were performed using default settings.

**Analysis of irradiation perturbed HSC datasets**

Quality control measures filtered cells based on total counts (<70000 and >5000), mitochondrial genes percent (<5%), ribosomal genes percent (<50% and >10%), and the number of expressed genes (>1500 and <7000). This resulted in 41252 cells. Genes expressed in fewer than 5 cells were excluded. Cells were normalized by total gene counts, and highly variable genes were identified using default parameters for PCA. Harmony data integration was employed using 100 components. The neighbor graph and Leiden clustering were conducted using default parameters, while UMAP was performed with a min_dist of 1.5. Cell types were annotated based on critical marker genes.

**Analysis of Dapp1 perturbed HSC datasets**

Cells underwent quality control, filtering based on total counts (<70000 and >1000), mitochondrial genes percent (<5%), ribosomal genes percent (<50% and >10%), and the number of expressed genes (>1000). This yielded 10224 cells. Genes expressed in fewer than 5 cells were excluded. Cells were normalized by total gene counts, and 2000 highly variable genes were chosen for downstream analysis. Data integration was performed using the Harmony algorithm. Default parameters were used for calculating the neighbor graph and UMAP. Leiden clustering was conducted with a resolution of 0.6, and cell types were annotated based on critical marker genes.
